# Zero-Shot Transfer of Protein Sequence Likelihood Models to Thermostability Prediction

**DOI:** 10.1101/2023.07.17.549396

**Authors:** Shawn Reeves, Subha Kalyaanamoorthy

**Affiliations:** University of Waterloo, Department of Chemistry, Waterloo, N2L 3G1, Canada

## Abstract

Protein sequence likelihood models (PSLMs) are an emerging class of self-supervised deep learning algorithms which learn distributions over amino acid identities in structural and evolutionary contexts. Recently, PSLMs have demonstrated impressive performance in predicting the relative fitness of variant sequences without any task-specific training. In this work, we comprehensively analyze the capacity of six PSLMs to predict experimental measurements of thermostability for variants of hundreds of heterogeneous proteins. We assess performance of PSLMs relative to state-of-the-art supervised models, highlight relative strengths and weaknesses, and examine the complementarity between these models. We focus our analyses on stability engineering applications, assessing which methods and combinations of methods can most consistently identify and prioritize mutations for experimental validation. Our results indicate that structure-based PSLMs have competitive performance with the best existing supervised methods and can augment the predictions of supervised methods by integrating insights from their disparate training objectives.

## Main

Enzymes are renewable catalysts capable of accelerating specific reactions under mild conditions, with wide applicability to industrial processes^1, 2^. Natural enzymes, however, evolve to operate within a tightly controlled cellular environment, and often cannot maintain their functional fold in the harsh and dynamic conditions required by some industrial reactions^3^. In general, modifications to the enzyme which stabilize its functional conformation relative to its unfolded state increase the longevity and cost-effectiveness of industrial enzymes, provided such modifications do not adversely affect activity, solubility, or other important attributes^1, 4^. A prevailing strategy to achieve this stabilization is to alter the protein sequence through mutagenesis. Although highly effective, brute force mutagenesis using techniques like directed evolution^5^ typically require the expression, purification and characterization of thousands of mutant enzymes before significant improvements are seen, due to the intrinsic inefficiency of introducing mutations without any bias toward those likely to be stabilizing^6, 7^.

Computational methods for mutant stability prediction can aid in the prioritization of promising mutants for experimental characterization. Most follow the same paradigm: a parametric model is developed to predict single-mutant stability from the wild-type sequence or structural features, the parameters of that model are tuned to minimize the error between predictions and experimental measurements of a stability-associated quantity, and that model is used to screen new candidate mutations for those which might be stabilizing. Typically, the stability measurement assesses the change in thermodynamic properties upon mutation; either the change in Gibbs free energy of unfolding (ΔΔ*G*) or the change in melting temperature (Δ*T_m_*). Biophysical methods, which comprise a trusted structure-dependent class of modelling approaches, are commonly used to predict the unfolding free energy change^8–12^. As a representative and popular example of biophysical modelling, Rosetta^8–10^ combines local conformational sampling for estimating the mutant’s native state with empirically-derived scoring functions for comparing the energies of the wild-type and predicted mutant structures, estimating the (ΔΔ*G*) as this difference. On the other hand, increased characterization of mutant proteins and advances in artificial intelligence methods have led to increasing use of machine learning for stability prediction. Typically, supervised machine learning methods are used to predict stability from some combination of the change in physicochemical properties of the wild-type and mutant residues, the features of the local chemical environment near the mutation site, and the relative likelihood of observing the wild-type and mutant residues in similar structural or evolutionary contexts^13–17^. The latter features are considered structural and evolutionary *statistical potentials*, respectively, and are known to be adequate stability predictors on their own^18, 19^. Foregoing conformational sampling, machine learning methods can make predictions orders of magnitude faster than biophysical methods, enabling the screening of many more candidate mutant enzymes for industrial applications. Notably, both biophysical and supervised machine learning methods are to various extents dependent on experimental stability measurements for fitting parameters, and both have been demonstrated to have significant shortfalls in terms of generalization^17, 20–22^.

The central problems limiting prediction accuracy relate to deficiencies in the training data: the current largest aggregate database for single-mutant ΔΔ*G* and Δ*T_m_*(FireProtDB) contains experimental observations for only a few thousand unique mutants across 237 proteins^3^. Taken as a whole, the existing data span a limited number of structural contexts, over-represent certain mutation types (such as those from alanine scanning experiments) while under-representing others (such as from one charged residue to another) and are heavily imbalanced in favor of the destabilizing class^10, 23^. Training data limitations have resulted in poor capitalization on advances in data-hungry deep learning, leading to stagnating improvement in prediction accuracy^17^. Furthermore, traditional models have limited practical applicability to stability engineering, since mutations predicted to be stabilizing have been shown to be neutral on average^20^, such that combining top-scoring mutations *in vitro* is unlikely to yield a significantly stabilized mutant. Consequently, only a limited number of examples of improved thermostability for industrial enzymes have been reported, and many of them involve Rosetta^24^. Until a plethora of additional stability measurements across protein families and mutation types become available, prediction of ΔΔ*G* or Δ*T_m_* must exploit more abundant data sources in order to fill the gaps. For example, Rosetta has already begun to include such auxiliary sources: the *BetaNov16* force-field was trained not only by optimizing rank correlation to experimental measurements from ProTherm^25, 26^ and from a semi-quantitative high-throughput *de novo* design study^27^, but also by optimizing native-sequence recovery and native-like amino acid distribution objectives.

Another emerging approach for overcoming these data limitations is through self-supervised pre-training of deep learning models. After being trained to predict partially masked-out residue identities using the remaining unmasked context in huge databases of protein structures, sequences, or sequence alignments, these *protein sequence likelihood models* (PSLMs) learn high-order and often long-distance interactions between residues which influence the distribution of evolutionarily viable amino acids in each biochemical context. Meier and colleagues recently demonstrated that a sequence-based model ensemble (ESM-1V) can make predictions of protein fitness based on the relative log-likelihood of mutant residues compared to the wild-type, with an average 0.509 Spearman correlation across 41 deep mutational scanning experiments^28^. This capacity for *zero-shot transfer* in which the unmodified pre-trained model is used to make predictions for a different task has also been observed in structure-based *inverse-folding* PSLMs. Specifically, multiple groups have reported success in zero-shot prediction of protease stability, binding affinity, and complex stability^29, 30^ while Dauparas and colleagues have already used generative modelling to design stabilized proteins and validate them *in vitro*^31^. Further discussion of PSLMs, including descriptions of those tested, is given in the Methods.

Given the impressive capacity of PSLMs to generalize from likelihood-based pre-training to protein engineering tasks, their performance in predicting thermostability relative to existing traditional methods is of immediate interest, particularly to protein engineers. With training objectives which are entirely different from previous methods, PSLMs have minimal risk of overfit or overestimation of test-set performance, which are pervasive problems in traditional modelling^17, 22, 23^. Additionally, they may be used in combination with existing models to augment predictions using complementary information. Finally, by modelling natural sequence likelihood, high-probability mutations are also expected to be enriched for frequently overlooked desirable attributes of industrial enzymes, such as enhanced function, solubility, and expression yield^1, 20^. In this study, we benchmark sequence likelihood models relative to existing supervised methods for protein stability prediction. We determine whether predictions are of comparable quality both in terms of classification and regression across a comprehensive dataset (FireProtDB^3^) and a held-out symmetric dataset (S461^23, 32^). We focus on stability engineering applications, especially the ability to correctly identify and prioritize stabilizing mutations for a variety of natural proteins, and perform a thorough analysis to understand the factors contributing to prediction quality and bias in PSLMs and other models. Finally, we show which behaviours, inductive biases and data sources might be complementary by comparing the performance of weighted combinations of all types of models and observing which ensembles lead to improved predictions, with consideration of the additional expense of computation time.

## Results

### Stability Prediction on FireProtDB

We studied six distinct approaches toward protein sequence likelihood modelling, coarsely divided into structure-based and sequence-based methods. To briefly describe their major differences, we assess three major structural methods: masked inverse-folding (MIF^30^), Protein Message Passing Neural Network (ProteinMPNN^31^), and evolutionary-scale modelling (ESM) inverse-folding (ESM-IF^29^), which have different architectures and are trained using monomeric structures from the Protein Data Bank, with the latter also using 12 million predicted structures from AlphaFold^33^. For sequence-based methods, ESM-1V is the only single-sequence predictor we used, trained using 98 million UniRef90 sequences, while Multiple Sequence Alignment Transformer (MSA-Transformer^34^) and Tranception^35^ are methods which also leverage sequence information from homologous proteins for inference, using very different strategies to do so. We elaborate greatly in Methods: Self-Supervised Learning. We first summarize the performance of PSLMs relative to a traditional biophysical method (Rosetta Cartesian DDG) and a traditional statistical potential based method (KORPM) on a lightly filtered version of FireProtDB, as shown in Table 1 (see Methods: Datasets). It is immediately clear that sequence-based PSLMs are less effective than other PSLMs and methods which incorporate structural information, a disparity which can be largely ascribed to the advantage that structure-based models have in terms of relevant information for evaluating biophysical interactions. However, the performance of structure-based PSLMs relative to traditional methods is less clearly distinguishable. When evaluated on the whole dataset, as is typically done, Rosetta Cartesian DDG (henceforth referred to as Rosetta) appears to be the superior model for predicting ΔΔ*G* in terms of binary classification (Matthew’s Correlation Coefficient, MCC), even when accounting for all possible classification thresholds (area under the precision-recall curve, AUPRC), and in terms of ranked regression (Spearman’s *ρ*). KORPM is also highly performant, scoring slightly above all PSLMs except in terms of AUPRC. When removing mutations which are neutral within experimental error (-1 kcal/mol < ΔΔ*G* < 1 or -2K < Δ*T_m_* < 2), classification performance increases across all models. Traditional methods seem to benefit from the removal of hard-to-classify mutations, but the PSLMs remain competitive (Supplementary Table 1).

**Table 1.** Comparison of Stability Models on the FireProtDB dataset. Sequence likelihood model performances are compared to traditional models and to random predictions, sorted by weighted *ρ* (w*ρ*), which cleanly divides all structural methods from sequence methods. The data are divided by measured quantity and by data aggregation method, with boldface values indicating the best performance in a given statistic and division. We drop 7 mutations for ΔΔ*G* and 2 for Δ*T_m_* which were not successfully predicted by KORPM. Italicized models (all sequence-based) had failed predictions for up to 14 unique mutations from the remaining 5058 (ΔΔ*G*) or up to 6 out of 2794 (Δ*T_m_*). Model subscripts indicate variants: training noise level for ProteinMPNN, monomer (M) for ESM-IF, 1 for a single subsampled MSA for MSA-Transformer, and 2 for model 2 of the 5-model ESM-1V ensemble.

| measurement | type | model | Whole-Dataset Statistics |  |  | Per-Protein Aggregate Statistics |  |  |
| --- | --- | --- | --- | --- | --- | --- | --- | --- |
| | | | MCC | AUPRC | $\rho$ | wNDCG | wAUPRC | $w\rho$ |
| $\Delta\Delta G$ | structural | Rosetta CartDDG | **0.326** | **0.533** | **0.602** | 0.908 | 0.615 | **0.421** |
|  |  | ProteinMPNN<sub>mean</sub> | 0.267 | 0.480 | 0.447 | **0.913** | **0.629** | 0.384 |
|  |  | ProteinMPNN<sub>30</sub> | 0.257 | 0.470 | 0.434 | 0.910 | 0.622 | 0.383 |
|  |  | ProteinMPNN<sub>20</sub> | 0.259 | 0.475 | 0.438 | **0.913** | 0.627 | 0.383 |
|  |  | ESM-IF | 0.211 | 0.443 | 0.374 | 0.906 | 0.595 | 0.380 |
|  |  | MIF | 0.211 | 0.441 | 0.433 | 0.909 | 0.591 | 0.377 |
|  |  | ESM-IF<sub>M</sub> | 0.203 | 0.437 | 0.364 | 0.905 | 0.584 | 0.362 |
|  |  | ProteinMPNN<sub>10</sub> | 0.243 | 0.466 | 0.433 | 0.908 | 0.614 | 0.357 |
|  |  | MIF-ST | 0.195 | 0.431 | 0.410 | 0.906 | 0.554 | 0.349 |
|  |  | KORPM | 0.275 | 0.466 | 0.498 | 0.898 | 0.568 | 0.337 |
|  | sequence(s) | *MSA-T<sub>mean</sub>* | 0.155 | 0.408 | 0.317 | 0.902 | 0.548 | 0.304 |
|  |  | *MSA-T<sub>1</sub>* | 0.138 | 0.395 | 0.307 | 0.898 | 0.538 | 0.281 |
|  |  | *ESM-1V<sub>mean</sub>* | 0.107 | 0.351 | 0.177 | 0.890 | 0.509 | 0.259 |
|  |  | *ESM-1V<sub>2</sub>* | 0.104 | 0.349 | 0.180 | 0.889 | 0.509 | 0.258 |
|  |  | *Tranception* | 0.097 | 0.341 | 0.237 | 0.891 | 0.506 | 0.254 |
|  | N/A | Gaussian noise | -0.005 | 0.276 | -0.005 | 0.844 | 0.420 | -0.005 |
| $\Delta T_m$ | structural | ESM-IF | 0.243 | 0.546 | **0.539** | **0.912** | 0.639 | **0.388** |
|  |  | ESM-IF<sub>M</sub> | 0.249 | 0.543 | 0.527 | 0.909 | 0.635 | 0.381 |
|  |  | Rosetta CartDDG | 0.249 | 0.501 | 0.510 | 0.906 | 0.645 | 0.362 |
|  |  | MIF | 0.253 | 0.527 | 0.453 | 0.907 | 0.642 | 0.350 |
|  |  | ProteinMPNN<sub>20</sub> | 0.311 | 0.566 | 0.506 | 0.905 | 0.668 | 0.347 |
|  |  | ProteinMPNN<sub>30</sub> | **0.316** | 0.568 | 0.498 | 0.905 | **0.676** | 0.335 |
|  |  | ProteinMPNN<sub>mean</sub> | 0.309 | **0.572** | 0.510 | 0.905 | 0.673 | 0.332 |
|  |  | ProteinMPNN<sub>10</sub> | 0.296 | 0.556 | 0.490 | 0.904 | 0.666 | 0.328 |
|  |  | MIF-ST | 0.206 | 0.498 | 0.409 | 0.900 | 0.627 | 0.316 |
|  |  | KORPM | 0.237 | 0.466 | 0.419 | 0.892 | 0.612 | 0.306 |
|  | sequence(s) | *ESM-1V<sub>mean</sub>* | 0.169 | 0.447 | 0.287 | 0.894 | 0.604 | 0.255 |
|  |  | *MSA-T<sub>mean</sub>* | 0.186 | 0.481 | 0.385 | 0.895 | 0.642 | 0.254 |
|  |  | *MSA-T<sub>1</sub>* | 0.187 | 0.485 | 0.378 | 0.894 | 0.642 | 0.232 |
|  |  | *ESM-1V<sub>2</sub>* | 0.151 | 0.444 | 0.299 | 0.890 | 0.601 | 0.228 |
|  |  | *Tranception* | 0.147 | 0.458 | 0.377 | 0.883 | 0.610 | 0.211 |
|  | N/A | Gaussian noise | 0.004 | 0.311 | 0.005 | 0.843 | 0.505 | -0.014 |

However, evaluating the statistics per-protein before aggregation changes the interpretation of the results significantly: compared to the default ProteinMPNN model, Rosetta is only superior in ranking the stability of all mutants (weighted-average Spearman’s *ρ*, w*ρ*; see Methods: Statistical Analysis), and by a narrow margin. When the task shifts to prioritizing stabilizing mutations (weighted-average Normalized Discounted Cumulative Gain, wNDCG) or discerning between stabilizing and destabilizing mutations (weighted-average area under the precision-recall curve, wAUPRC), 3 of 4 ProteinMPNN variants are superior. This observation suggests that Rosetta’s performance is inflated by correctly discerning between destabilizing mutants more than stabilizing ones, an idea supported by later findings. While a bootstrapped analysis (see Results: Model Complementarity) shows that the improved performance of ProteinMPNN relative to Rosetta depends on the selection of data, aggregating per-protein results produces a more logical comparison where Rosetta is no longer clearly superior to inverse-folding methods. Meanwhile, for Δ*T_m_* prediction, all structural (inverse-folding) PSLMs are strongly competitive with Rosetta regardless of the class of statistic, with ESM-IF scoring a significantly higher w*ρ* and wNDCG, while MIF scores also higher wNDCG and ProteinMPNN (trained with 0.30 Å backbone noise) scores the highest wAUPRC. Further, all inverse-folding models score higher than the simpler statistical potential-derived supervised method, KORPM, across evaluations for the Δ*T_m_* subset.

The training objective of both traditional supervised methods (and most other existing methods) relies at least in part on minimizing the error between predictions and experimental observations of ΔΔ*G* across mutants from many protein families pooled together, where the available training data consists mostly of destabilizing mutations. Hence, it is not surprising that these two methods perform better on whole-dataset metrics considering mostly destabilizing mutations, and on ΔΔ*G* measurements. However, this training objective is at odds with the goal of stability engineering, which focuses on the correct relative ranking and prioritization of stabilizing mutations for a single protein at a time. As such, test statistics like root mean squared error and accuracy are less meaningful for this task, while statistics about pooled proteins can confound results by comparing mutants of unrelated proteins. Rosetta’s pooled-protein ranking (rather than error) objective and auxiliary sequence recovery and native amino acid distribution objectives may be contributing factors in its retained per-protein performance relative to KORPM.

Unlike PSLMs, which have never been trained on experimental stability measurements, Rosetta and KORPM are trained on a large fraction of FireProtDB^10, 23^, and interesting behaviors arise when holding out a major constituent of their training data, the ProTherm dataset (see Supplementary Table 2). While all methods maintain roughly the same performance on the reduced ΔΔ*G* set (1677 unique mutations) indicating good generalization, performance of supervised methods on the reduced Δ*T_m_*dataset (1006 unique mutations) drops drastically: both methods now perform similarly to sequence-based methods in terms of per-protein aggregate statistics. Meanwhile, PSLM performance changes less appreciably. However, most of the observations remaining after removing measurements belonging to the ProTherm dataset originate from ThermoFluor characterization of protein complexes (including 6G4B, 1PX0, and 6TQ3) which relies on the binding of dye to exposed hydrophobic patches and may simply indicate partial dissociation of monomers rather than unfolding or inactivation. However, this phenomenon may still influence fitness, since the structure-based PSLMs retain more of their predictive power. Rosetta’s training objective attempts to optimize many objectives simultaneously while adjusting relatively few parameters, which may limit its flexibility to model more complex fitness phenomena, perhaps explaining its loss of performance here. Supplementary Figure 1 compares the ranking performance of Rosetta Cartesian DDG and ProteinMPNN across individual proteins and helps to illustrate these unusual cases.

### Mutant Feature Analysis

We next seek to identify the features of mutations which are predicted with variable success by different models, focusing on the full ΔΔ*G* set to mitigate confounding factors and provide a conservative estimate of the relative performance of PSLMs. Additionally, we use the weighted Spearman’s *ρ* statistic to most equitably compare between methods with different underlying distributions. We discover two major discriminants of model performance between model types, illustrated in Figure 1. The first discriminant is the stability itself: when FireProtDB ΔΔ*G* measurements are grouped into discrete categories, major limitations become apparent. First, Figure 1(a) shows that all methods struggle to rank each protein’s stabilizing mutations: sequence-based models make predictions which are worse than chance, while all other models make predictions only slightly better than chance. Meanwhile, evolutionary models have the best performance when ranking significantly destabilizing mutations. Since these mutations are significant enough to be deleterious to fitness and such mutations comprise a significant fraction of the landscape in deep mutational scans, this observation is consistent with the established performance of evolutionary methods in predicting variant fitness^28, 35^. It should be noted that increased stabilization of already-stable proteins seldom confers a significant fitness advantage, which explains the incapacity of evolution-conditioned models to rank stabilizing mutations. This is further corroborated by Figure 1(b), which shows that evolutionary models do not predict stabilizing mutations as being more likely than moderately destabilizing ones. Notably, there may be a trade-off in the capacity to rank stabilizing and destabilizing mutations for PSLMs which depends on degree of evolutionary character: ESM-IF and MIF-ST exhibit similar patterns to the sequence-based models. MIF-ST uses sequence transfer (ST) to infuse information from sequence-based embeddings into the structure-based graph representation, while ESM-IF trains using millions of structures generated by AlphaFold, which is an evolution-conditioned structure predictor^33^.

**Figure 1.**
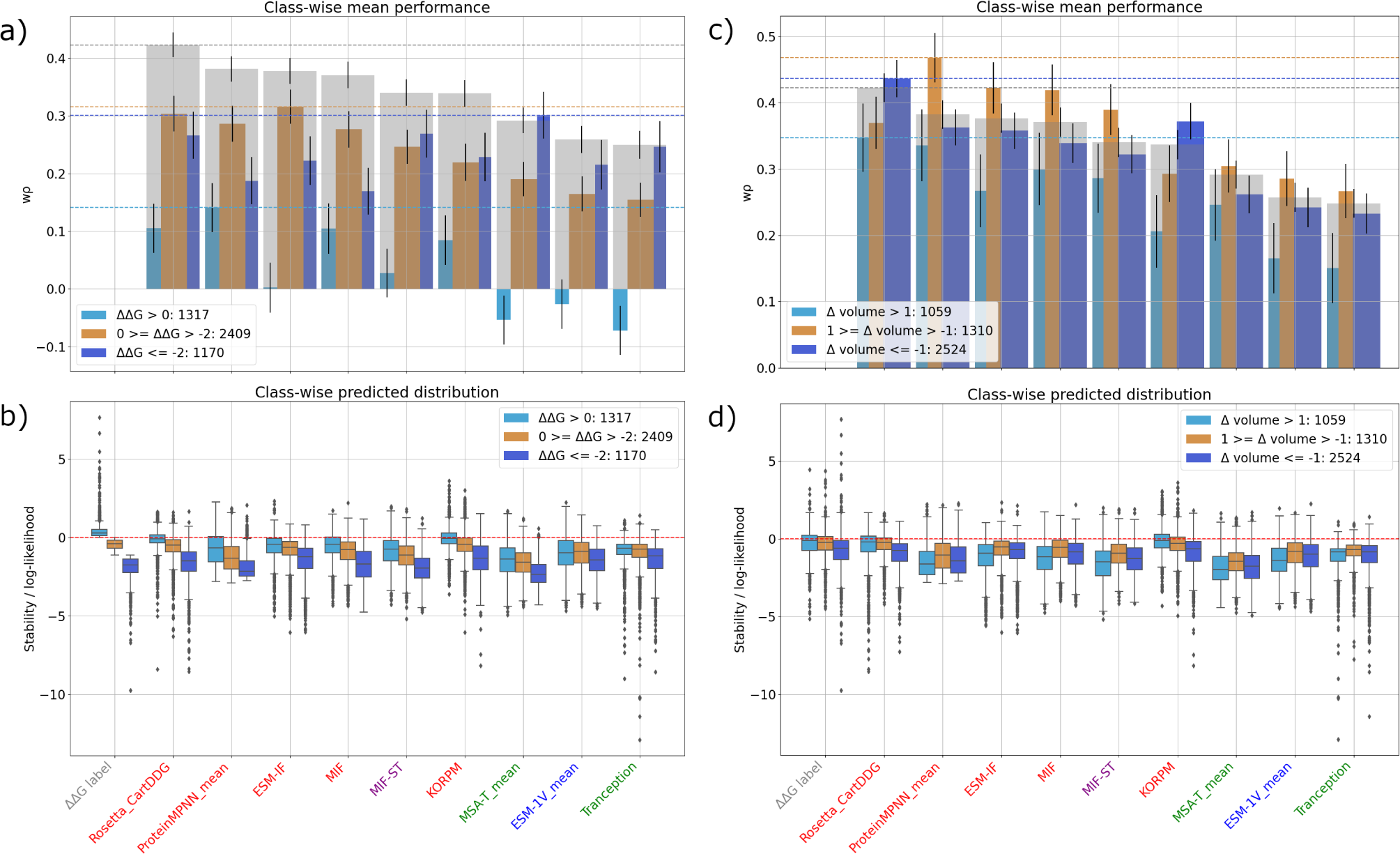
Performance Breakdown based on Mutational Features. Plots (a & c) illustrate the change in weighted Spearman’s *ρ* (w*ρ*) when grouping the dataset based on stability (a) or change in volume upon mutation (c), versus the ungrouped data (transparent grey) across 1000 bootstrapped replicates (error bars). Horizontal dashed lines on the two plots indicate the best performing model’s average score for each group. The items are sorted with the highest ungrouped performance at the left, and bar widths indicate the number of mutants contributing to that statistic, which is also shown in the legend. Variation in the total number of mutants occurs mainly due to excluding data slices when there were less than 2 proteins to rank for a given protein and feature range. Plots (b & d) indicate the distribution over all predictions of the same groups as (a & c), with the ground truth label distribution on the far left. The red horizontal dashed line indicates the decision threshold for mutations considered to be predicted stabilizing, which is used for MCC and classification metrics. X-axis labels are color-coded by model type: grey for labels, red for structural, blue for sequence-only, purple for structural and sequence features, and green for evolutionary models.

Three physicochemical features of mutations with particular relevance to stability are the solvent-accessible surface area, change in hydrophobicity, and change in volume upon mutation. Indeed, these features alone can be used for effective supervised modelling^17, 36^. Through binning mutations on the basis of volume change calculated using the Quantiprot Python package^37^ in Figure 1(c), we note that PSLMs consistently rank mutations with minimal volume changes more effectively than those with volume decreases and much more effectively than those with significant increases. Inverse-folding models learn from static structures: unlike Rosetta, they cannot accommodate bulkier substitutions through rotamer sampling or local backbone rearrangements. Hence, as shown in Figure 1(d), volume increasing substitutions are predicted as the least likely class; the models cannot be expected to correctly rank these mutations when many are anticipated to cause steric clashes, which would not be observed frequently in the training data. However, ProteinMPNN uses a robust training strategy in which backbone noise is added to the structures, which may generate spurious clashes and account for its improved performance over other inverse-folding methods. On the same line of reasoning, inverse-folding models appear to be superior predictors of similar-volume mutations, likely because of their improved generalization capacity acquired from heterogeneous training structures. Rosetta’s conformational sampling strategy may allow it to compensate for volume-shrinking mutations more effectively than volume-increasing ones, which can cause more widespread structural rearrangement. Unusually, bulky substitutions are the most stabilizing class in this dataset and have a near-neutral median stability; supervised models appear to have captured this bias, while the PSLMs indicate that these tend to be less likely in natural contexts.

We account for the other two relevant physicochemical features in Supplementary Figure 2, but do not find major determinants of PSLM versus traditional model performance. Instead, we observe an increase in ranking capacity of all models for buried mutations to more hydrophilic residues, perhaps because such interactions are the most specific and hence provide all models with predictive signal. As known zero- and few-shot predictors of structure, even sequence-based models show good capacity to differentiate such interactions. We also find that supervised methods predict hydrophobic substitutions as more likely to be stabilizing, mostly in line with the ground truth distributions. Meanwhile, PSLMs show a distributional shift to lower probability for hydrophobic surface mutations relative to hydrophilic ones. The predicted infrequency of such mutations reflects their consequences to fitness: hydrophobic surface mutations can promote misfolding and aggregation, impair function, and reduce solubility. They tend to increase ΔΔ*G* by enabling the burial of additional hydrophobic surface area relative to the unfolded protein, but this behavior is also likely to disrupt the functional fold^38^. Hence, this learned bias may lead to undesirable effects. We also show, in Supplementary Figure 3, that PSLMs (and especially those with higher evolutionary character) have diminished performance for scoring mutations when the same residue in homologous sequences is highly conserved. This observation is consistent with the training procedures, and the improved performance of inverse-folding methods in the high-conservation case suggests that these methods are better able to generalize from non-homologous structures. However, most inverse-folding methods still perform significantly worse than the traditional methods for conserved residues. We also show in Supplementary Figure 4 that, as one of only two models which consider quaternary structure explicitly in the input (the other being ProteinMPNN), ESM-IF shows the best performance in ranking mutations of multimeric structures. The effect of including the full biological assembly is made especially clear through comparison to the monomeric version (ESM-IF(M)), which shows significantly diminished performance for multimeric structures only. However, the performance is not significantly different from Rosetta, which only receives the monomeric structure.

### Evaluating PSLMs for Stability Engineering Objectives

We next investigate the screening utility of PSLMs for engineering applications, where many candidate mutations are screened and a limited number of top mutations are selected for experimental validation. We first assess the precise detection of stabilizing mutations at various percentiles of ranked model predictions (Figure 2, a & b). When considering ΔΔ*G* (a), ProteinMPNN has increasingly greater performance than Rosetta beyond the 70th percentile: if the screening budget is anywhere between the top 1% and top 30% of predicted stabilizing mutations of a given protein, ProteinMPNN would be expected to yield the highest fraction of experimentally stabilizing mutations. Other inverse-folding methods perform similarly, but do not consistently outperform Rosetta, as the summary statistics in Supplementary Table 3 indicate. In the case of Δ*T_m_* (b), ProteinMPNN consistently and significantly outperforms Rosetta, with the latter hovering around the problematic sticking point of 50% stabilization above the 75th percentile^20^. It is also interesting to note that for both measurements, MSA-Transformer (the best performing sequence-based method for this task) performs poorly except for the top few percentiles, perhaps because a small subset of FireProtDB comprises variants with an increased prevalence in homologs due to improved stability, and hence, high likelihood according to MSA-Transformer. For similar reasons, MSA-Transformer achieves the highest mean stability for predicted stabilizing mutations (relative log-likelihood > 0, shown in Supplementary Table 3) for ΔΔ*G*, although this is partly attributed to its high specificity (and hence, limited recall).

**Figure 2.**
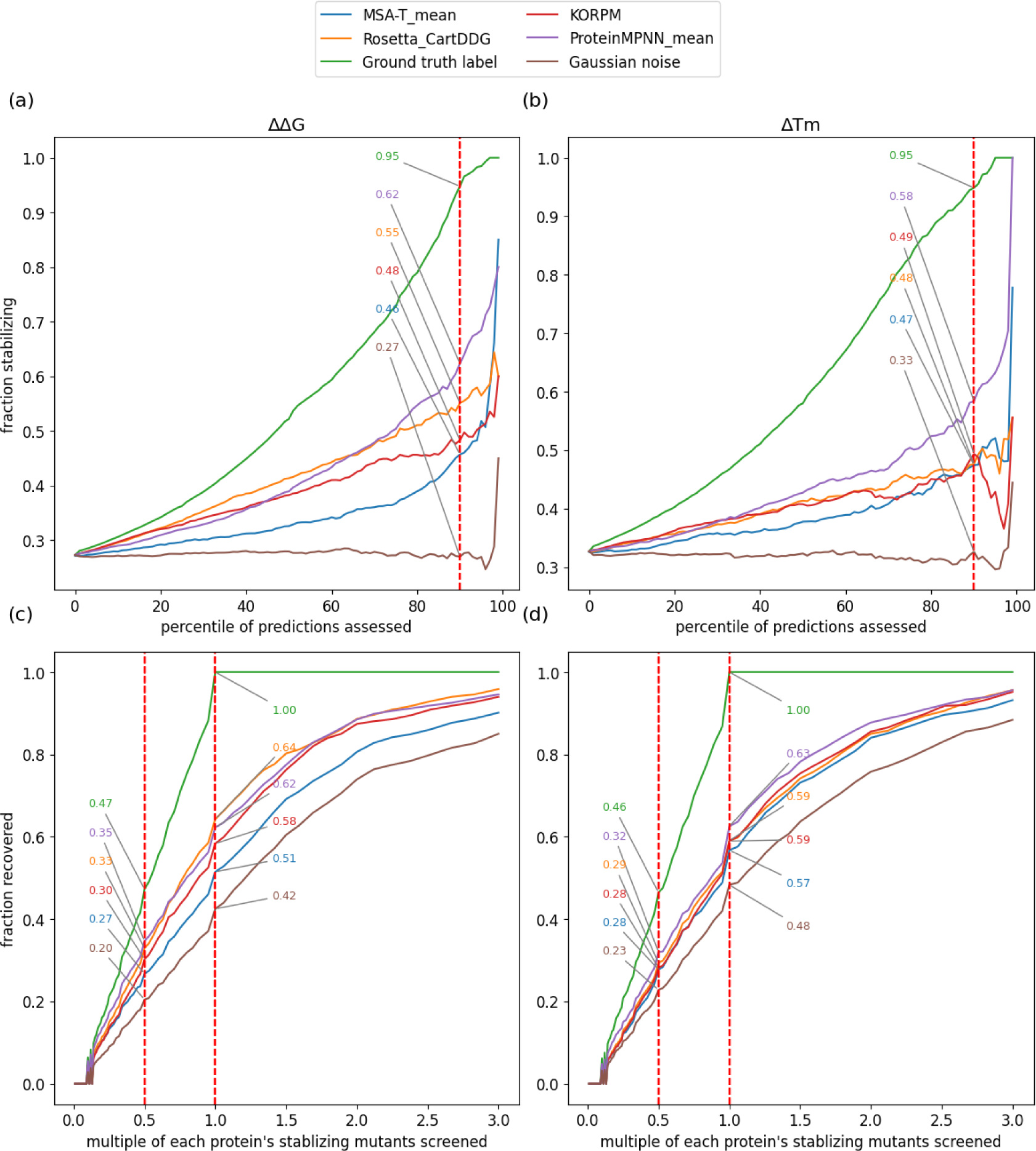
Visualization of Stable Mutant Enrichment in FireProtDB. The top plots (a & b) show a percentile-wise precision curve (PPC): for a given percentile of top-scoring mutants according to the models, what fraction are experimentally stabilizing? Percentiles are identified per protein and the results are aggregated across all proteins; a protein with only 5 characterized mutants will contribute to the visualization only in discrete steps of 20%. There is a high amount of variance beyond the 95th percentile as few proteins have more than 20 mutations, and predictions are only rendered for experimentally characterized mutants. The vertical dashed line indicates a plausible screening budget in which the 90th percentile of mutations (top 10% scoring) is experimentally validated, at which precision is annotated. In the lower plots (c & d), the recovery (recall) of all mutations with positive stability measurements is assessed with a screening budget of various multiples *K* of the number of experimentally stabilizing mutations per protein (K-times recovery curve, KXRC). There is not exactly 50% recovery at 0.5x for the ground truth labels due to rounding-down odd numbers of stabilizing mutations. Annotations of exact scores for 0.5x and 1.0x screening budgets are recorded.

In Figure 2, c & d, we show how many of a protein’s top predictions would need to be screened for each before a fraction of the total stabilizing (ΔΔ*G* or Δ*T_m_* > 0) mutants are recovered. While the former curves consider precision, these curves are focused on recall. ProteinMPNN recovers more stabilizing mutations than Rosetta across almost all screening budgets for Δ*T_m_* and until 70% of the number of true stabilizing mutations are screened for ΔΔ*G*, with similar performance thereafter. Notably, no method recovers 80% of stabilizing mutations until at least 1.5x the number of stabilizing mutations is screened, and less than 96% of stabilizing mutations are recovered at 3x. We summarize the area under both curves for all models as well as many other stability engineering focused statistics in Supplementary Table 3. While KORPM and Rosetta have more true positive predictions than PSLMs, they also predict more false positives, and the average stabilizing effect of predicted positives is very close to zero.

### Model Complementarity

We investigate the performance of weighted pairs of models to explore model complementarity and whether additional computation can justify potential performance gains. Specifically, we tested predictive performance when adding together predictions from distinct models, each normalized by their individual standard deviation across the whole dataset. We tested combinations of the six PSLM models and their variants (see Methods: Self-Supervised Learning), as well as Rosetta Cartesian DDG and KORPM. Additionally, we incorporate the three simple features mentioned previously as a minimal predictive set: change in volume, change in hydrophobicity, and solvent accessible surface area of the wild-type residue. In each pair, one model has weight equal to 1, and the other model can have weight equal to 1, 0.5, or 0.2 We test only 3 weights to restrict the ability to fit the dataset and thus to report generally applicable results, and hence obtain 5 unique combinations of weights per pair, for a total of 525 combinations. It bears repeating that Rosetta and KORPM are not blind to this dataset, and hence results involving these models may be over-optimistic. However, the focus of this section is on the augmentation conferred by predictions which are made in a zero-shot fashion, and hence, performance *improvements* are expected to be generalizable.

Figure 3 reports the screening performance and runtimes of the best performing pair from each set of binary combinations, as well as the statistical significance of the improvement, if any. We choose to report the cumulative stabilizing effect if applying all predicted stabilizing mutations to concretely illustrate the utility of screening mutations *in silico*. This visualization reveals that Rosetta can benefit significantly by infusing statistical information from all PSLMs, with all but Tranception resulting in a greater net stabilization than when combining KORPM with Rosetta instead. The best combination applies an equal weight of 1 to ProteinMPNN (with a training backbone noise of 20 Å) and greatly improves upon Rosetta’s performance (P=10*^−^*^48^, paired t-test, 100 bootstrapped replicates), with negligible additional time-cost relative to the expensive biophysical sampling procedure. However, it should be noted that the total stabilizing effect is still small relative to the potential net stability of all stabilizing mutations: the Rosetta + ProteinMPNN combination achieves roughly 18% of the (mean bootstrapped) net stability of 832 kcal/mol. We report Rosetta’s total runtime as though predictions were run in series (summing each mutation’s runtime) rather than in parallel (which would consider only the maximum runtime). This estimate is appropriate when using a consumer workstation (with one GPU and one CPU) rather than a compute cluster. Additionally, it allows for direct comparison to other methods (which were all computed in series, but on a GPU for all PSLMs) and accounts for the total computational expense. In Supplementary Figure 5, we breakdown the runtimes into individual predictions to more clearly illustrate the magnitude of disparities between methods.

**Figure 3.**
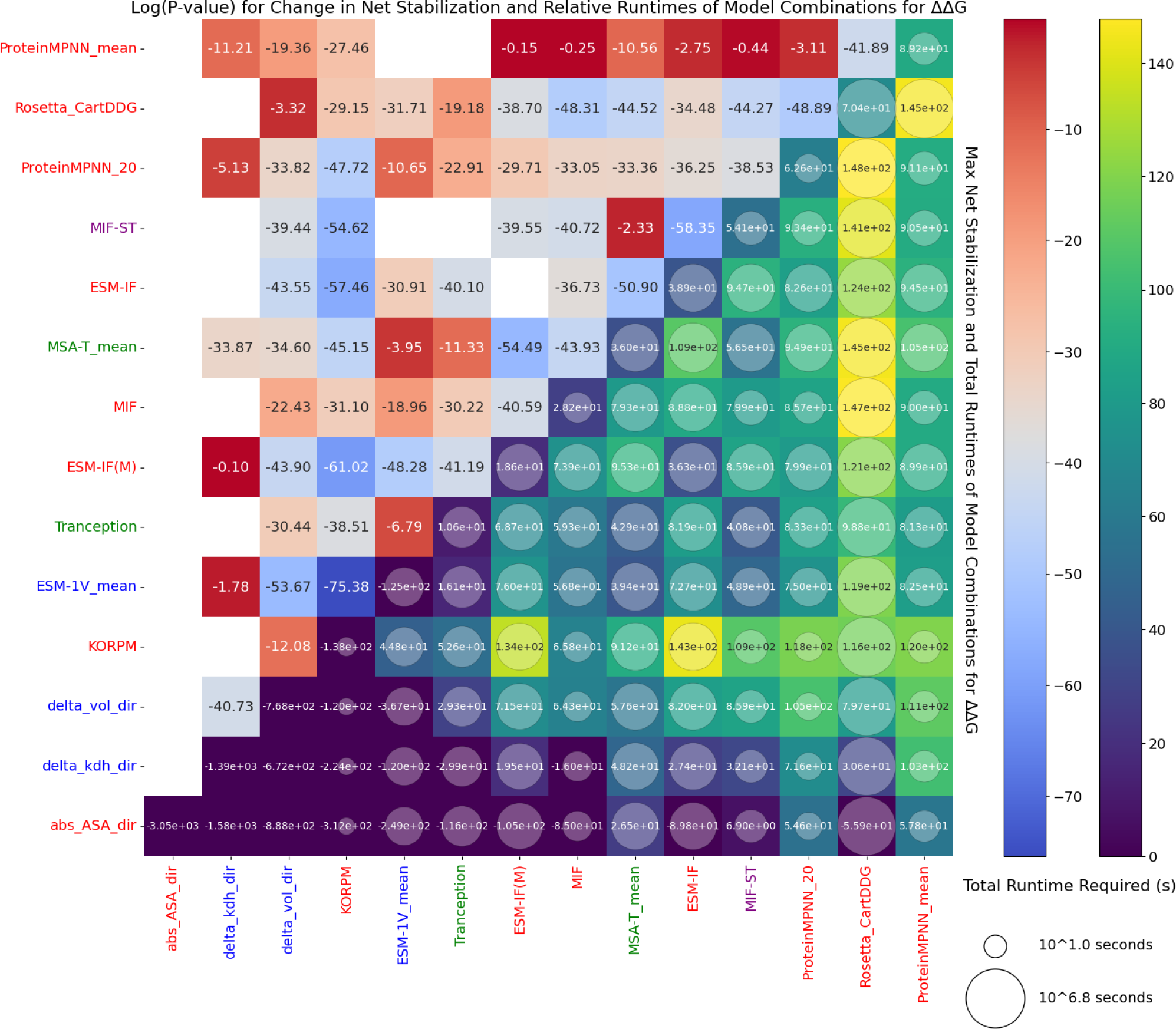
Screening Performance and Runtimes of 2-Model Combinations on FireProtDB. Colors and annotations in the triangle on the lower right indicate the net stabilizing effect of applying all predicted stabilizing mutations to all proteins (in kcal/mol) for the best of five weighted combinations of normalized predictions of two models. Performance of individual models is given on the diagonal. Combinations and models which do not have net-positive stabilization are colored purple. The sizes of the circular markers’ radii are proportional to the log(runtime) for making both sets of predictions. The annotations and colors on the upper left of the plot indicate the log(P-value) for improvement in the net stabilizing effect for the best combination, compared to the best constituent model (paired t-test, 100 bootstrapped replicates). White cells indicate that performance of all combinations was reduced.

Many combinations of models lead to meaningful performance gains. Indeed, yellow cells in the lower triangle indicate that most combinations of traditional methods with PSLMs are among the best performing, with the combination of ProteinMPNN with KORPM compromising between performance and speed and exceeding Rosetta’s performance while predicting all mutants in just over 2 minutes. Importantly, the superior performance when combining traditional models and PSLMs is seen whether examining ΔΔ*G* or Δ*T_m_*, using other statistics including whole-dataset AUPRC and per-protein weighted Spearman’s *ρ* or wNDCG (Supplementary Figures 6-8), and even across datasets, as we discuss later. We find that the combination of Cartesian DDG + ProteinMPNN(mean) * 0.5 leads to the most consistent high performance across metrics. The upper-left triangle of the plots indicate that combining traditional and PSLM methods usually results in more highly significant performance gains (over the best-performing constituent model) than combining PSLMs. Interestingly, PSLMs benefit by incorporating the change in volume to augment predictions for classification (supplementary Figure 6) and ranking (Supplementary Figure 7) tasks, suggesting that it may be possible to compensate the deficiencies seen in Figure 1. Similarly, incorporating the change in hydrophobicity improves ranking and prioritization performance (Supplementary Figures 7 & 8) significantly. The same effects are absent or less extreme for traditional models, suggesting the latter have already been adequately calibrated to these features. Incorporating volume and polarity change into PSLM predictions likely counterbalances the bulky- and hydrophobic-substitution favoring test set noted previously. Despite their high performance, PSLMs are not optimized for stability prediction out-of-the-box and may have room for significant improvement with supervised fine-tuning or through using their predictions as features for supervised models. Finally, we find that although all models except KORPM achieve positive mean stabilization, the high specificity of MSA-Transformer (seen in Supplementary Table 3) can be used to achieve combinations with the highest degree of stabilization per predicted-positive mutation (Supplementary Figure 9), suggesting that MSA-Transformer combinations may be useful in the low-throughput *in vitro* regime.

### Symmetric Held-Out Data

We study the performance of PSLMs and a collection of high-performing traditional methods, mostly falling under the umbrella of supervised approaches, on the S461 dataset^23^ which includes 41 proteins each with no more than 25% sequence similarity to any data used for training supervised models. Modelled structures of mutants were provided by the authors of the original S669 superset^16^, augmenting the dataset with an equal number of stabilizing mutations and enabling model antisymmetry and bias to be quantified. Additionally, predictions for all mutants in the parent S669 set were rendered using numerous modern methods^16^, which we also compare to PSLMs here. More details are given in Methods: Datasets.

We begin by considering only the direct mutations, since these are more applicable to the engineering objective: statistics about predictions of hypothetical inverse mutations can be dominated by highly-stabilizing reversion mutations from deleterious variants which are not expected to occur frequently in natural proteins or engineering targets. Figure 4 shows the net stability enhancement from combinations of two models under identical conditions to Figure 3 but shows the Spearman correlation between individual models’ predictions in the upper triangle rather than the improvement log(P-value). Unlike in FireProtDB, the only single model to achieve net positive stability is Tranception, indicating the increased difficulty of this benchmark in terms of stabilizing mutations. Indeed, S461 is a much smaller dataset than FireProtDB, with a distributional shift toward destabilizing mutations (Supplementary Figure 10), meaning there is much less potential stability gain possible. The feature distributions also differ somewhat, shifting toward volume shrinking (direct) mutations, with more solvent exposure, and higher wild-type conservation on average. Based on the previous analyses, these characteristics are expected to favor traditional methods.

**Figure 4.**
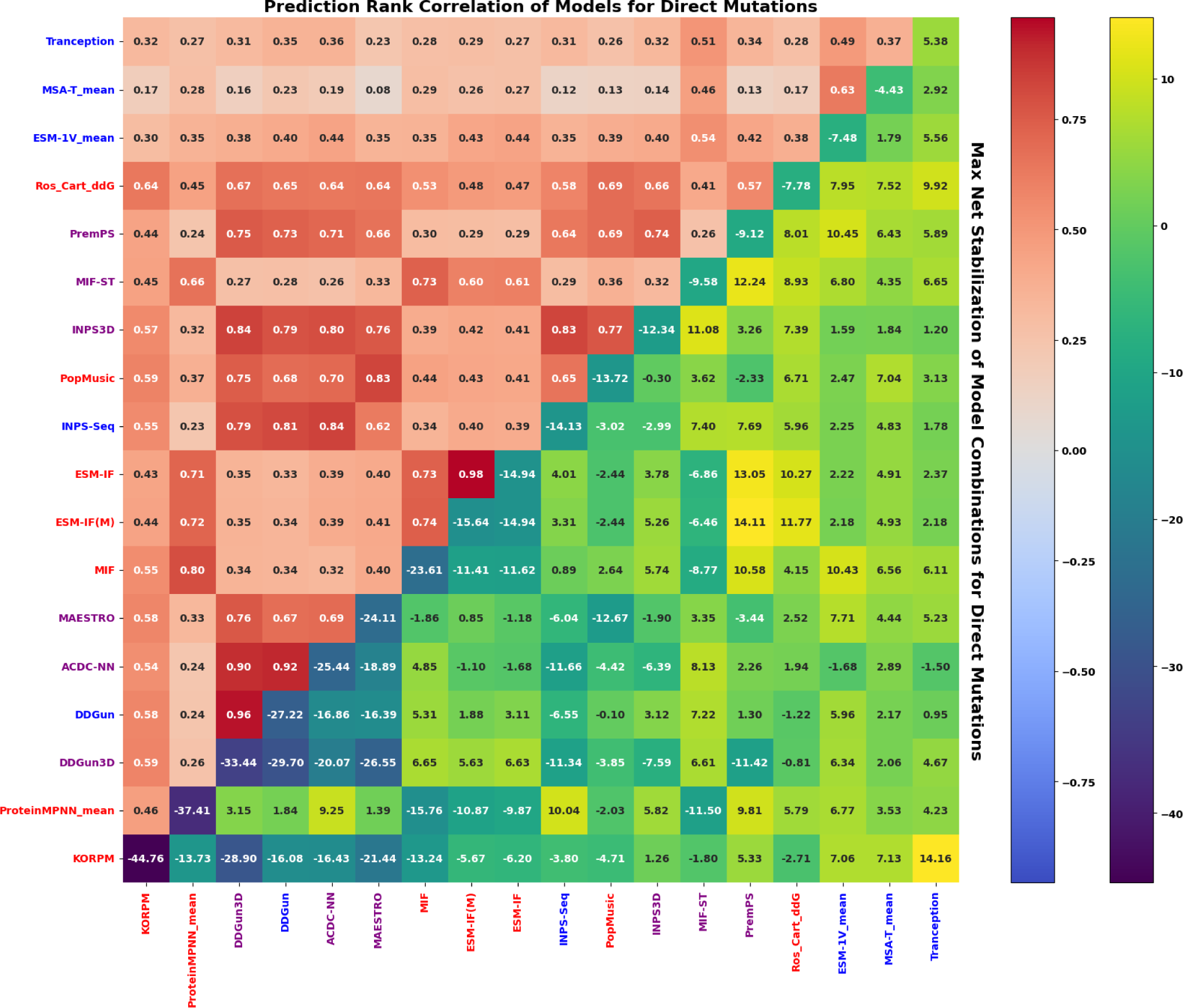
Screening Performance and Correlations of 2-Model Combinations on S461. The annotations and colors in the upper left of the plot indicate the Spearman correlation between all predictions of high-performing models. Performance of individual models in terms of combined stabilizing effect of all predicted-stabilizing mutations (in kcal/mol) is given on the diagonal, while the lower triangle indicates net stabilization for the best of five weighted combinations of normalized predictions of two models (no bootstrapping). Only monomeric structures are available for the inverse mutations, so only monomeric variants of ESM-IF are shown.

While the performance of inverse-folding models is diminished, sequence-based PSLMs surprisingly come out ahead. The substantial positive net stability achieved by many combinations is promising, as past works have noted that predicted mutations tend to be experimentally neutral on average, such that adding together many predicted positive mutations (assuming independent effects) is unlikely to result in significantly enhanced stability^20^. We also note an apparent symmetry across the diagonal of Figure 4, where less correlated individual predictors tend to achieve higher net stabilization when combined, suggesting that the performance gains from ensembles arise through combining disparate insights, ultimately arising from different inductive biases, training data, and training objectives. We assemble Supplementary Table 4 by merging the top 10 unique combinations of models according to weighted NDCG, area under the percentile-wise precision curve, and net stabilization into a single table. While no combination consistently performs best, 23/29 combinations involve one traditional model and one PSLM, and the equally-weighted combination of KORPM and Tranception is a top performer in terms of wNDCG and Net Stabilization, hence why fewer than 30 combinations are listed. Additionally, most combinations achieve a significant fraction of the potential stability gains, which amount to only 37 kcal/mol in the best case for this dataset.

Next, we consider the more general case of the bidirectional set (both direct and reverse mutations assessed together), which contains a total of 922 unique mutations and can potentially better represent situations where numerous stabilizing mutations are possible, probing the versatility of model combinations. As shown in Supplementary Figure 11, models which make less biased predictions tend to rank mutant stability for bidirectional mutations more accurately than biased combinations, such as combinations of inverse-folding models and any of MAESTRO, Rosetta, PopMusic, INPS-3D, and other inverse-folding models. We repeat the same selection of top models as for the direct mutations and compare them in Supplementary Table 5, finding that these comprise only 22 unique models comprising exclusively combinations involving at least one PSLM, where the second model always has very high antisymmetry, and may also be a PSLM. Notably, Rosetta is entirely absent, partly due to its destabilizing bias. Whether considering direct or bidirectional predictions, and regardless of the choice of statistic, the most predictive ensemble pairs usually combine one supervised method with a PSLM.

Finally, we demonstrate that a pair ensemble selected based on its performance on the FireProtDB can greatly enhance the screening of stable mutations for unseen data. For the best weighted pair on FireProtDB (Rosetta Cartesian DDG + ProteinMPNN * 0.5) according to consensus between net stabilization (Figure 3), AUPRC, weighted NDCG and weighted Spearman *ρ* Supplementary Figures 6-8), we assess its performance for protein engineering objectives relative to the best-performing individual models tested on the S461 set. We return our focus to the prediction of direct mutations for previously stated reasons. Despite its modest performance in terms of Net Stabilization (Figure 4), this combination is one of the top-10 ranked combinations in terms of area under the percentile-wise precision curve (Supplementary Table 4). As shown in Figure 5, this combination is superior to all other individual methods, including its constituents, particularly in the prediction of direct stabilizing mutations above the 90th percentile, which represents an intuitive screening budget. Similarly, it shows superior recovery of stabilizing mutation for most screening budgets less than or equal to the true number of mutations. In the case of inverse or bidirectional mutation sets, the precision and recall curves are again improved over the constituent methods (Supplementary Figure 12). While the improvements in each case are modest, that consistent improvements in the detection of stabilizing mutations can be made to an already high-performing method across datasets simply by infusing protein likelihood information suggests the potential of these methods to greatly enhance predictions with further engineering.

**Figure 5.**
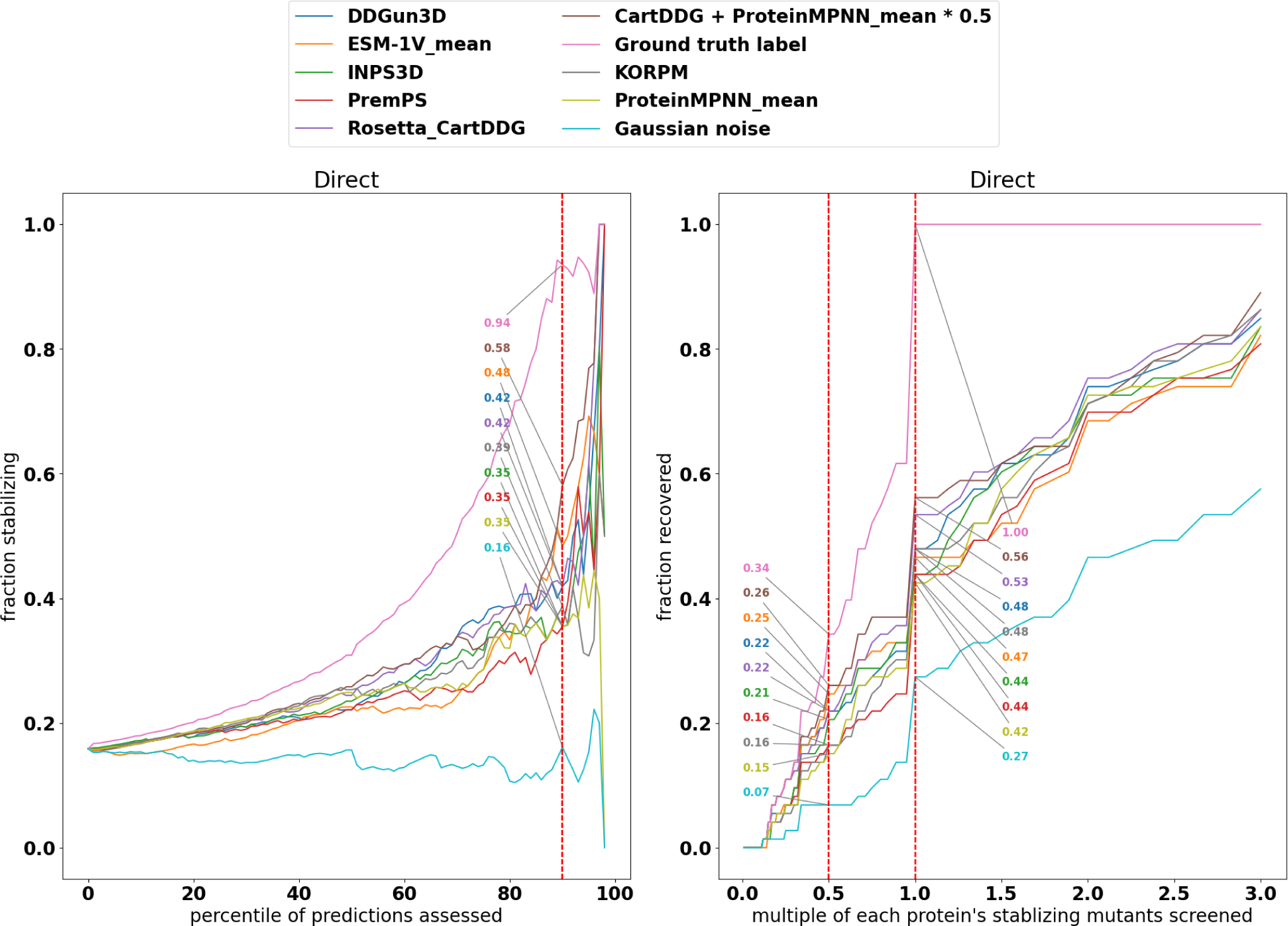
Visualization of Stable Mutant Enrichment by Pair Ensemble in S461. The left plot shows percentile-wise precision: for a given percentile of top scoring mutants according to the models, what fraction are truly stabilizing? Percentiles are identified per protein and the results are aggregated across all proteins. The vertical dashed line indicates a plausible screening budget in which the 90th percentile of mutations (top 10% scoring) is screened. In the right plot, the recovery (recall) of all mutations with positive stability measurements is assessed with a screening budget of various fractions of experimentally characterized stabilizing mutations per protein.

## Discussion

The accurate prediction of protein stability upon mutation is a challenging endeavor, but can greatly accelerate protein engineering efforts through the prioritization of stabilizing mutations. We demonstrated one avenue to push through the stagnating performance of stability prediction methods: the incorporation of information about the likelihood of observing a sequence in nature, especially given its structure, as modelled by state-of-the-art deep learning approaches. We suggested that such models, which we refer to as PSLMs, have a complementary set of strength and weaknesses compared to traditional methods, which depend on experimental stability data. PSLMs and traditional methods have a similar ability to rank, classify and prioritize mutations among the majority of quantitative single-mutant stability measurements recorded so far, but when examining the performance of each method on feature-based subsets of data, traditional methods and PSLMs have clear strengths relative to one another: PSLMs perform poorly when there is a substantial change in residue volume upon mutation and when the mutated position is highly conserved in homologs, while traditional methods including Rosetta and KORPM have learned biases such as predicting surface mutations to hydrophobic residues as stabilizing, despite the consequences this may have to other protein engineering objectives like soluble expression. Additionally, ProteinMPNN appears to have superior performance in ranking and prioritizing stabilizing mutations in particular, while evolutionary and sequence-only PSLMs are best at ranking significantly destabilizing mutations with relevance to predicting loss-of-function variants and understanding disease. The complementary strengths and separate data sources are two contributing factors toward the consistently higher prediction and screening performance seen in combinations of traditional models and PSLMs.

As deep mutational scanning experiments allow for an ever-increasing capacity to generate new (semi-quantitative) stability-associated measurements for training and testing models, and new approaches for high-throughput expression of tailored mutants becomes increasingly accessible, the role of expensive biophysical techniques for engineering stable mutations may be increasingly relegated to domains such as refinement and analysis of pre-screened candidates. For instance, new experiments^39^ are achieving semi-quantitative characterization of nearly one million mutations, a domain in which validating Rosetta would be computationally intractable. Meanwhile, deep learning methods can make rapid predictions for testing or screening at scale and can benefit hugely from the emerging plethora of data, perhaps by fine-tuning of PSLMs which clearly have useful prior assumptions for this task. In this work, we demonstrate that PSLMs innately predict mutant stability surprisingly well. Since PSLMs can greatly benefit from crudely combining their knowledge with existing supervised methods, or sometimes even with rudimentary physicochemical features, it is almost certain that fine-tuning with potentially millions of data points drawn from a variety of proteins would further enhance performance, although this may come with impacts on function and solubility which are hard to predict..

## Methods

### Self-Supervised Learning

We explore likelihood-based protein stability prediction using six distinct self-supervised model types and some of their variants that employ slightly different inputs or modelling approaches. We categorize these models by the type of information used for training and inference: (i) sequence-only models, including ESM-1V, which depend solely on the primary amino acid sequence of the wild-type protein; (ii) evolutionary models, including Tranception and MSA-Transformer, which depend on a multiple sequence alignment of homologous protein sequences, and lastly (iii) inverse-folding models, including ESM-IF, MIF(-ST) and ProteinMPNN, which utilize the knowledge of the three-dimensional structural fold and therefore require an experimental or predicted structure of the wild-type protein of interest. Owing to their similar architecture, behavior and inputs, we frequently group the first two categories into “sequence-based” models and contrast them with “structure-based” inverse-folding models. All PSLM models considered here are trained in a purely self-supervised way, using a variant of the masked language modelling objective. In this procedure, a model predicts a fraction of the residues which have been hidden (masked) or mutated (corrupted) based on the remaining information^40^. The model is expected to learn to predict the actual identity of the masked or corrupted tokens, which is accomplished by tuning the parameters to minimize a loss function that compares the predicted distribution to the known identity of the predicted residues (cross-entropy loss).

#### Sequence-Based Models

Sequence-only and evolutionary (multiple sequence alignment-based) self-supervised methods use variants of the Transformer architecture, which has seen extraordinary success in the field of language modelling^41^. Such models do not use any structural information as input, instead encoding sequences or sequence alignments using sets of tokens (one token per residue) with positional embeddings added to each token to allow the model to reason about the position of a residue in a sequence, and potentially to which aligned sequence it belongs, as in the MSA Transformer^34^.

The defining feature of the transformer architecture is the self-attention mechanism, which allows the model to learn how to produce output representations from input tokens based on long-range contextual information. For a given protein sequence, a key, query and value are extracted from each token using learnable projection matrices. For each input token in the sequence, a new output representation is generated from a convex linear combination (the “attention” distribution) of values associated with the keys of other tokens, where the weights are assigned according to the similarity of the input token’s query representation to each other token’s key representations, including the input key. Hence, the model can infuse information from other positions relevant for understanding the context of each token, improving the information-density of its updated representation. To adapt the self-attention strategy to multiple sequence alignments, the MSA-Transformer implements two separate attention distributions: a row attention shared by all sequences, which determines which other parts of its sequence a residue position can attend to, and a column attention, which determines which tokens in the same aligned position from sequences can be used for context. Intriguingly, Rao and colleagues showed that the row attention distribution predicted by these models is correlated with the contact map for the corresponding protein structure, enabling state-of-the-art zero-shot contact prediction based on the extraction of co-evolutionary relationships^34^. The self-attention procedure is repeated across multiple attention heads with their own projection matrices, resulting in different notions of contextual encoding. The representations from the different heads are linearly combined according to another learned set of parameters.

Following the self-attention algorithm, a single learnable feed-forward neural network is applied to each output token representation, giving the model more flexibility to encode non-linear relationships between the features of each token embedding. The multi-head self-attention, position-wise feed-forward neural network, and additional residual connections and layer normalizations form a block which can be stacked upon other blocks to produce deeper representations. High-level representations following numerous encoder blocks have been an area of intense focus in the field of *protein language modelling*, as they have been shown to be information-dense inputs for downstream tasks such as functional annotation^42, 43^, structure prediction^34, 42, 44^, and even thermostability prediction via transfer learning^44–46^. For masked language modelling (and likelihood inference), following the final encoder blocks, a linear layer is applied to each token embedding to produce a distribution over possible residues at that position. During training, only the masked positions are used to calculate the loss to the ground-truth label (the correct residue).

In this study, we explore only ESM-1V in terms of sequence-only models, as this was the only sequence model we found trained specifically for variant-effect prediction in a purely self-supervised manner^28^. In terms of evolutionary methods, we examine MSA-Transformer^34^ and Tranception^35^. Rather than stacked encoder blocks culminating in a discriminative final layer, Tranception uses stacked decoder blocks and autoregressive (generative) decoding, like the generative pretrained transformer (GPT) architecture of OpenAI^47^. Autoregressive models provide the flexibility to iteratively generate a new sequence, conditioning new residue predictions on all previous predictions and provided residue context. Furthermore, Tranception implements retrieval to augment new token predictions with the empirical distribution at the aligned position, hence using an abstraction of the MSA compared to MSA-Transformer, and blending in some sequence-only character through its training on unaligned sequences^35^. A comparison of these models is given in the Table 2 below.

**Table 2.** Summary of Models.

| Model | Params | Architecture | Inference | Training Data | Masking |
| --- | --- | --- | --- | --- | --- |
| MSA Transformer | 100M | 12 Transformer encoder layers with axial attention and tied row attention | all-at-once | 26M MSAs with an average of 1192 sequences each | uniform 15% masking over full MSA |
| Tranception | 700M | 36 Transformer decoder layers with varied-size depth-convolutions in attention heads | bidirectional autoregressive with retrieval | 250M UniRef100 sequences with retrieval of related sequences | random sequence mirroring, forward and reverse averaged in testing |
| ESM-1V | 652.4M | 33 Transformer encoder layers, ensemble of 5 models | all-at-once | 113M UniRef90 2020-03 cluster representative sequences | 15% corrupted by (80% mask, 10% mutation, 10% unchanged, with loss calculation) |

The multiple sequence alignment exploited by evolutionary models (MSA-Transformer and Tranception) is generated after searching sequence databases such as UniRef100^48^ using tools such as JackHMMER^49^, which finds homologs (related protein sequences) iteratively using Hidden Markov Models. The limited inductive biases of the Transformer architecture used by all sequence-based models necessitates a large corpus of training data; for instance, 26 million MSAs averaging 1192 sequences each were used to train the 100 million-parameter MSA-Transformer^34^. Evolutionary models can be used in a “few-shot” transfer learning context: that is, they are provided with positive examples of natural sequences (the MSA) to shape their inference of new sequence likelihoods. In contrast, sequence-only ESM-1V is evaluated in a zero-shot context: it neither receives training specific to the new task (stability prediction, in this case), nor any additional context outside the remainder of the predicted sequence. With 650 million trainable parameters, the model stores evolutionary information in its parameters rather than extracting it in inference; ESM-1V used 98 million diverse sequences derived from the UniRef90 2020-03 database to establish these relationships^28^.

#### Structure-Based Models

Inverse-folding models learn to predict the likelihoods of amino acids using the backbone atom coordinates for a structure of interest (among various other structural features, such as dihedral angles), as well as the identities of amino acids at positions which are not being predicted, and therefore rely on sequence information as well. Their name is derived by contrast to “forward-folding” structure prediction models like AlphaFold^33^: rather than predicting the folded structure of a protein given its sequence, inverse-folding models predict the sequence from the folded structure.

Inverse-folding models mostly make use of the graph neural network (GNN) architecture, in which each residue is represented as a node, and spatially proximal nodes are connected with edges, using the folded structure to determine the spatial arrangement of the graph^50^. More specifically, ESM-IF^29^, MIF^30^ and ProteinMPNN^31^ all use variants of the message-passing neural network (MPNN) architecture for encoding structural information. Briefly, each node contains quantitative information about the residue (particularly relating to the spatial relationships between its atoms and neighboring residues), and information can be propagated to connected nodes to generate updated node representations with contextual information about the local environment. In this way, it is similar to the Transformer, but the contextual information is more spatially local. In MIF-ST, *sequence transfer* is used to encode residue-level information from additional sequence-based Transformer embeddings from CARP in the nodes^44^; for this reason, we sometimes refer in figures to MIF-ST as a *hybrid* between sequence and structural methods. Edges are undirected and contain additional attributes (such as the distance between the *C_β_* atoms of the two nodes it connects), and the concatenation of features from all nodes and edges connected to a given node are transformed using a neural network, forming a “message”, which is propagated across all connected edges. All three models have edges between a constant number of nearest neighbors. Upon aggregating incoming messages from all connected nodes and edges, the node representation is updated using this message aggregate according to a second neural network. The three models differ slightly in terms of the order in which edge and node updates occur and which attributes are input to the neural network, and ESM-IF uses a geometric vector perceptron in place of the MPNN. For each architecture, the same neural network parameters are adaptable to arbitrarily many structures (they exhibit combinatorial generalization), as the same neural networks are reused for information propagation throughout all connections and all input structures^50^. Hence, the GNN architecture is parameter efficient relative to the Transformer architecture.

Graph neural networks are capable of impressive sequence recovery in the context of sequence likelihood modelling. For instance, ProteinMPNN was shown to recover 52.4% of the residues in held-out native sequences, which is notable considering the non-optimality of native sequences for their backbones and the limited restrictions on surface residue identities^51^. The strong inductive bias of locality and high sample efficiency of inverse-folding models compensates their limited training data: inverse-folding models are largely limited in terms of training to the number of experimentally determined three dimensional structures of proteins available in the Protein Data Bank or redundancy-reduced datasets from CATH, but these training datasets can be augmented with predicted structures generated with algorithms such as AlphaFold^29, 33^.

While each inverse-folding model encodes similar input structural information in a similar way, their decoding strategies (which decide the relative probabilities of different residues in context) vary significantly. MIF and MIF-ST use a variant of the BERT encoding scheme^40^, resulting in embeddings which directly model the residue probabilities based on the entire masked input sequence at once. Meanwhile, like Tranception, ESM-IF and ProteinMPNN use autoregressive decoding. ESM-IF uses the transformer architecture for left-to-right sequence decoding, with the limited inductive bias offset by the larger training dataset augmented by 12 million AlphaFold-predicted structures^33^. Meanwhile, ProteinMPNN uses the most flexible decoding scheme (order agnostic decoding), allowing for sequence context from all other residues (not just those upstream in sequence as for ESM-IF) to inform the identity of a single-site variant.

In this study, we explore ESM-IF^29^ (based on full oligomeric structures or target chain only and designated with the M suffix, for monomer), MIF^30^ (and variant MIF-ST, which also uses sequence embeddings as additional node-level input), and ProteinMPNN^31^ (with variants trained with 0.1, 0.2 or 0.3Å added backbone noise and the ensemble mean of these 3; e.g. suffix 10 = 0.1Å). A comparison of these models is given in Table 3 below:

**Table 3.** Summary of Models

| Model | Params | Encoder | Decoder | Inputs | Inference | Training Data |
| --- | --- | --- | --- | --- | --- | --- |
| ESM-IF | 142M | 4-layer<br>GVP-GNN<br>+ 8-block<br>Transformer | 9-block<br>Transformer | N, Ca, C coordinates | left-to-right<br>autoregres-<br>sive | 16,153 single-<br>chain CATH<br>v4.3 structures +<br>12M AlphaFold<br>structures |
| Protein-MPNN | 1.678M | 3-layer<br>MPNN | 3-layer<br>MPNN | distances between N,<br>Ca, C, O, and virtual<br>Cb, primary<br>sequence distance,<br>chain identity | order-<br>agnostic<br>autoregres-<br>sive | 23,358 PDB<br>assemblies <30%<br>identity, <3.5<br>Angstrom resolu-<br>tion |
| MIF(-ST) | 3.4M (+<br>640M for<br>MIF-ST) | 4-layer<br>MPNN | None (linear<br>on nodes) | dihedral and planar<br>angles, Cb atom<br>distances, (corrupted)<br>sequence | all-at-once | ~19,000 CATH<br>v4.2 structures (+<br>42M UniRef50 se-<br>quences) |

### Prediction Methodology

#### Protein Sequence Likelihood Models

The information which can be provided as input to each method (even among inverse-folding models) for inference is different. ProteinMPNN and ESM-IF receive the most complete set of information, including the sequence and structure (atomic backbone coordinates) of all chains in the biological assembly, and ProteinMPNN receives two additional atom coordinates over ESM-IF as shown in the above table. Meanwhile, MIF and MIF-ST receive only the target chain’s sequence and structure. All sequence methods (MSA-Transformer, Tranception, and ESM-1V) recieve the full UniProt sequence of only the target chain for inference, meaning they also have access to information unknown to the inverse-folding methods, which may be especially relevant when the experimentally solved structure is severely truncated. Finally, MSA-Transformer and Tranception also receive redundancy-reduced, high-coverage sequences obtained using homology-searching of UniRef100 with the UniProt sequence as the seed (see Input Data Generation and Preprocessing). Because Tranception uses a retrieval scheme, the MSA ends up being reduced to the distribution of residues at each position, meaning some information is lost.

The basis and calculation of predictions of single-substitution mutations from each model depends on its inference scheme. For MSA Transformer, ESM-1V, and MIF(-ST) the inference scheme is all-at-once, meaning that the entire sequence (with the mutated position masked) is passed as an input to the model simultaneously, with the identity of all other positions being used for inference. In addition, for MIF(-ST), the geometric information for the entire protein, including the masked residue, is also provided. These models generate probability distributions over amino acid identities at each position which is evaluated using a heuristic that compares the log-probability of the mutation against the wild-type residue at the position of interest Eqn. 1. By treating the difference in log-probabilities as a stability prediction, we assess the model’s performance in predicting the specific mutations in a zero-shot manner; a difference greater than zero suggests that the mutant residue is more likely than the wild-type, and hence is potentially more stable.

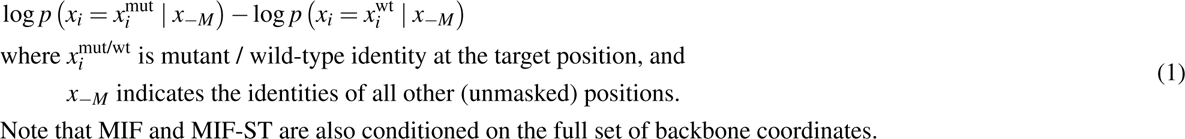

In contrast to the all-at-once nature of the other models, Tranception, ESM-IF, and ProteinMPNN are autoregressive models: the sequence is decoded token-by-token in a generative manner, and the probability distribution at a mutated position of interest depends on the identities of only the previously decoded sequence (and either the retrieved distribution at the aligned position or the encoded structure context, as described previously). Since the remainder of the mutated sequence is pre-determined for single substitutions (i.e., the task is not generative), we feed the largest possible window of existing sequence to the decoder as context according to the protocols described in the Input Data Generation and Preprocessing subsection. For these autoregressive models, we compare the log-probabilities of the full structure-derived sequence, meaning that the full sequence must be decoded for the joint probabilities of the mutant and wild-type to be compared. The evaluation proceeds according to Eqn. 2.

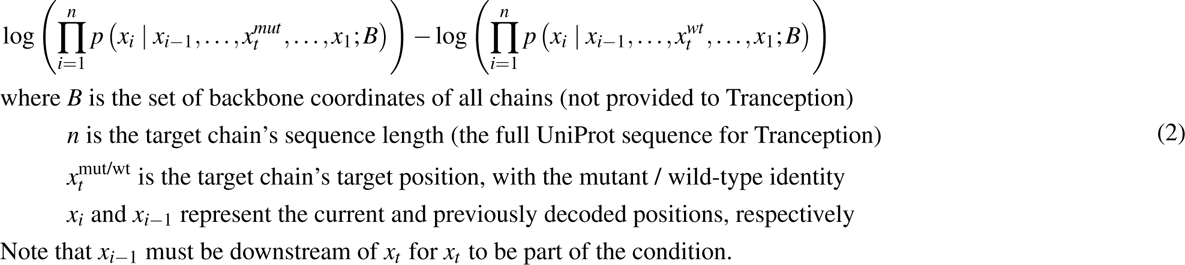

Unless otherwise stated, we use zero as the decision boundary for classification purposes, such that a mutation is predicted to be stabilizing only if the log-likelihood difference is greater than zero, i.e. the mutant sequence is more likely than the wild-type. We always use only the wild-type structure for inference, except for computing inverse mutations, in which the (predicted) mutant structure^16^ is considered to be the “wild-type”. In the case of ProteinMPNN(mean) we report the mean of predictions made by models trained with three different levels of added backbone noise. ESM-1V is shipped as an ensemble of 5 models, from which we also report the mean unless otherwise stated. Finally, MSA-Transformer was previously shown to generate superior fitness predictions when aggregating outputs from 5 subsamples of a filtered alignment (as explained in the Data Generation and Preprocessing section), so we follow this procedure and report the mean of the 5 predictions unless otherwise stated. For some of the above models, particularly ProteinMPNN, multiple choices of additional hyperparameters or preprocessing decisions are possible. We reproduce original works as closely as possible and report default models in the main text. Inference steps were performed on a single A100 GPU for all PSLMs.

#### Biophysical Modelling

We choose Rosetta as a primary methodology to benchmark sequence likelihood models against. Rosetta comprises a trusted toolkit for protein engineering and design, and stability measurements can be made rigorously using a combination of conformational sampling and scoring using a semi-empirical forcefield^9, 10, 26^. At the cost of high computational expense, Rosetta offers a consistently high standard for single-substitution mutant stability prediction across numerous studies, although the parameters used to make predictions can vary^23, 45, 46^. We compared two versions of Rosetta: a “traditional” version of Rosetta stability prediction, which uses the recommended parameters for the ddg_monomer application according to Kellog et al.^26^ and a “best-known” configuration for the cartesian_ddg application^9, 10^ reproducing the parameters and procedure used by Hoie et al^19^. However, we found that ddg_monomer almost always gave inferior performance, and often failed to render predictions in an acceptable amount of time, so we excluded it from the analysis. Notably, the two version methods use different forcefields for scoring, with the cartesian_ddg protocol using the more recent beta_nov16_cart and each performing different extents and types of conformational sampling. In both cases, PDB structures were first relaxed according to their corresponding protocols. The exact methodology used is given in the accompanying repository (see Data Availability. Predictions were generated using parallel runs, each on a single CPU, as the program is designed to run. We used Rosetta version 2019.21.60746.

#### Statistical Potential

We consider KORPM, a relatively simple structure-based statistical potential based on relative orientations of residues, recently shown to have state-of-the-art performance on the held-out Ssym dataset^23^. KORPM is related to inverse-folding models, in that both are derived from the relative probability of observing a mutant versus wild-type residue in a structural context. However, KORPM decomposes the statistical potential into contributions between all residues in contact with the native residue and applies learned weights to each contribution based on amino acid type. While its simplicity and strong natural inductive bias (the statistical likelihood) mitigate overfit, it is difficult to establish whether its performance is generalizable to all types of mutations, and as a new method, has remained to be externally validated. Predictions were generated in series on a single CPU, and runtimes were estimated by dividing the whole-dataset runtime by the number of mutations.

#### Other Methods

In this work, we refer to both modelling approaches above as supervised methods, since they use parameters which are fit to experimental stability measurements. Meanwhile, DDGun^52^ selects simple weights based on correlations of individual features to training data (a very weak form of supervision) while methods like SDM^18^ have no learnable parameters. We group these methods and supervised methods under the more general umbrella of *traditional methods*. All other methods appearing in the results are either supervised methods (e.g. PopMusic^53^, INPS^14^, PremPS^13^, and MAESTRO^15^) or physicochemical descriptors (Δ volume, Δ hydrophobicity, and SASA).

### Datasets

Authors of SLM methods variously report the zero/few-shot prediction of stability of de novo designed mini-proteins^29, 30^ from Rocklin et al.^27^, complex stability for binding proteins from SKEMPI^29, 30^ according to Townshend et al.^54^, and indirect observations of stability from deep mutational scanning experiments^28, 35^. However, the performance of state-of-the-art sequence likelihood models for mutant stability prediction of a wider range of structurally diverse natural enzymes remains to be performed. Additionally, an understanding of the characteristics of mutations which are more accurately predicted or are predicted as more stabilizing by different methods is desired, to better match the choice of model(s) to the specific task at hand or use them in complementary ways. For instance, models biased toward predicting surface mutations to hydrophobic residues may not be well-suited toward engineering proteins which are only marginally soluble. In this study, we address these knowledge gaps and report the performance of sequence likelihood models on two datasets: FireProtDB and S461.

#### FireprotDB

FireProtDB^3^ is an aggregate dataset containing most of the published direct measurements of single-substitution effects on thermostability on a wide variety of proteins. As such, it also recapitulates the biases toward certain types of mutations which are present in the original datasets it aggregates. Measurements are split into ΔΔ*G* and Δ*T_m_*, with many independent experimental observations of some proteins, but just one observation for others. We filter entries with no indicated structure and those whose structure does not contain the mutated part of the sequence, and use the first reported structure for entries with more than one listed. Also, we take the median over mutations with multiple reported values for experimental measurements of the same type in order to reduce the effect of erroneous measurements with large magnitude. We use the full dataset, including both values labelled as curated and not curated. We separate the analyses of ΔΔ*G* and Δ*T_m_*, and a summary of the filtered FireProtDB data used in our analyses is given in Table 4.

**Table 4.** Summary of Filtered Measurements from FireProtDB. Note that the sum of the two measurement types is not the same as the combined value, since there are mutations whose ΔΔ*G* and Δ*T_m_* are both recorded.

| Measurement | Observation | Unique muts | Stabilizing | Structures | Muts / Struct | wt->Ala |
| --- | --- | --- | --- | --- | --- | --- |
| $\Delta\Delta G$ | 38760 | 5065 | 1381 | 176 | 1-937 | 1358 |
| $\Delta T_m$ | 14191 | 2794 | 913 | 131 | 1-308 | 608 |
| Combined | 52923 | 6313 | 1918 | 230 | 1-937 | 1525 |
| Curated | 34138 | 5614 | 1688 | 191 | 1-932 | 1525 |

#### S461

S461^23^ is derived from S669^55^, which is a symmetric dataset, meaning it contains complementary experimental measurements and (modelled) mutant structures for hypothetical reversions of each mutation. It is also held-out from most supervised methods including Rosetta, KORPM, and all other methods tested. Sequences in this dataset have no more than 25% similarity to any sequence within the most popular training dataset, S2648^56^. Structures for all 669 mutants are generated using Robetta by the original authors, allowing for the comparison of the prediction of structural methods for mutations and their corresponding reversions, which should have equal magnitude and opposite sign. We also use the predictions rendered from various (mostly supervised) stability prediction methods reported by the Pancotti et al. in parts of our analysis^16^. Notably, this dataset is not perfectly orthogonal to FireProtDB, but this is inconsequential for our purposes. We used the S461 dataset, a further curated subset of S669 produced by the authors of the KORPM method^23^. This curated data corrects some erroneous records in the original dataset and eliminates additional mutations likely required for natural function, such as those at oligomer interfaces. It contains experimental structures for 48 wild-type proteins, with 1-68 mutations per protein and total of 461 mutations each with a single experimental ΔΔ*G* measurement.

#### Input Data Generation and Preprocessing

To facilitate data comparison, we adopt the sign convention for ΔΔ*G* of *unfolding*, such that proteins stabilized by mutations have both positive ΔΔ*G* and Δ*T_m_*. This is a notable departure from the more commonly used ΔΔ*G* of *folding*, in which stabilizing mutants have negative sign; we make this choice to simplify comparisons, particularly with Δ*T_m_*. Unless otherwise stated, we use zero as the threshold above which all experimental measurements of mutations are considered stabilizing for classification purposes.

We obtain biological assemblies for each wild-type structure in either dataset, and for NMR-resolved structures, the first model from the ensemble is used. To maximize the compatibility of sequences and structures between models, sepharose (SEP), L-3-aminosuccinimide (SNN) and acetyl (ACE) pseudo-residues are deleted while all other non-canonical residues except selenomethionine are converted to UNK. Missing residues are modelled using the complete_pdb function from Modeller^57^. The structures are then passed to each model leveraging structural information, which may perform additional internal preprocessing. Oligomeric proteins were prepared according to model requirements and abilities. For ProteinMPNN, positions from monomers sharing identical sequences are tied for inference, meaning the geometric average of the residue probability distributions across identical chains is used. In the case of MIF and MIF-ST, where handling of oligomeric structures is not implemented, and ESM-IF(M), since assessing both monomeric and multimeric predictions is recommended, only the mutated chain is used for inference.

The residue mutations from different models that utilize different inputs are compared by mapping the sequence indices between Uniprot and the PDB structures. The mutation index from the FireProtDB is based on UniProt numbering. We use the pairwise global alignment algorithm (pairwise.globalms) available in the BioPython library for mapping the structure to the Uniprot sequence. Manual checks and mapping are done for specific cases, such as models with missing residues. Conversely, S669 provides the mutation indices in structural terms, which are mapped back to the sequence obtained from the PDB structure using the PDB e-KB^58^. In cases where the UniProt sequence is not available, the PDB structure’s sequence was used in its place as input for sequence-based models. The full UniProt sequence is provided as context for inference using ESM-1V when it is less than 1022 amino acids in length (2 tokens are reserved for the start and end of the sequence). When the sequence is longer, we first capture the whole sequence of the associated structure for comparison with structural methods (always less than 1022 residues), then extend backward toward the N-terminus, and finally extend past the end of the structure if additional tokens remain (e.g. the structure was truncated relative to the UniProt sequence). Further, ESM-1V is an ensemble of 5 models trained with different seeds; we experiment with using the mean and median of all 5 predictions for the zero-shot stability estimate and report the (marginally) better-performing mean.

To generate the multiple sequence alignments (MSAs) used in Tranception and MSA Transformer, we use JackHMMER to search UniRef100 (obtained July 2022) using the full UniProt sequence as a seed. Specifically, we search for 8 iterations using a bitscore threshold of sequence length divided by 2. If fewer than 400 sequences are retrieved, the process is repeated once with a new bitscore threshold of sequence length divided by 4. Next, the hhfilter function from HHSuite^59^ is used to enforce a minimum of 75% coverage and 90% identity between sequence in the alignment. The resulting alignment passed directly to Tranception, or further subsampled 5 times for MSA-Transformer (with replacement between iterations only) according to the same sequence weights used in traditional reweighting^60^, unless it contains fewer than 384 sequences, in which case the entire MSA is passed. Hence, most predictions made by MSA-Transformer are aggregated from 5 predictions with different prompts, according to the mean (the median was again slighly worse). We use the same context window designation for MSA-Transformer as for ESM-1v. In the case of Tranception, up to 1023 tokens can be fed as previous context, and an additional 1023 tokens can be added following the mutation to augment the right-to-left stage of the decoding. We follow the original procedure, using the default implementation to provide the maximum available context.

Finally, in the case of ProteinMPNN, there is a flexible decoding order. Consequently, we choose the mutated residue to be decoded last such that it receives information about all other residues, including geometric information about the target position, to infer the identity distribution. There is a small amount of stochasticity in predictions related to the decoding order of the remaining residues.

### Statistical Analysis

We focus on statistics with the most relevance to protein engineering applications, prioritizing those which treat class-imbalance and non-linear relationships appropriately, and facilitate the understanding of performance on stabilizing mutations in particular. An important aspect of this analysis is the treatment of datasets both as a whole and through aggregated assessments made per-protein. Per-protein statistics are important for estimating performance when engineering a single protein, e.g., when screening and then characterizing a limited number of mutations, and we use various aggregation strategies to meaningfully represent generalization capacity.

#### Whole-Dataset Statistics

Matthews Correlation Coefficient (MCC) is a statistic used to evaluate the quality of binary classification models; hence, it requires binarization of both predictions and labels, which occurs at the default threshold of 0. It is a robust metric that considers true positive (TP), true negative (TN), false positive (FP), and false negative (FN) predictions, where the positive class is the stabilizing mutations. The coefficient ranges from -1 to 1, where 1 indicates a perfect prediction, 0 signifies a random prediction, and -1 indicates a completely inverse prediction. MCC is highly regarded for its ability to account for imbalanced datasets and to deliver a balanced assessment of a classifier’s performance, regardless of the class sizes. However, it is only defined when there are examples of each class and predictions of each class, and hence it cannot be used per-protein, partly since not all proteins have stabilizing mutations. The formula for calculating the Matthews Correlation Coefficient is:

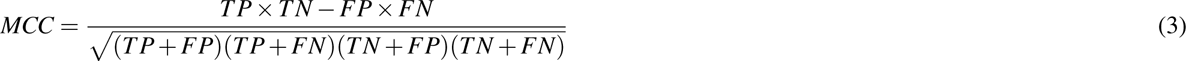

Area under the precision recall curve (AUPRC) is a summary statistic of the average performance of a binary classifier across different decision thresholds. Precision is the probability that a mutation is stabilizing, given that it is predicted as such: TP / (TP + FP). Precision can be traded off for improved recall (also referred to as sensitivity), which is the fraction of stabilized proteins detected: TP / (TP + FN). AUPRC considers all decision thresholds for the distinction between stabilizing and destabilizing predictions, which is useful since the threshold parameter may be optimized beyond the intuitive threshold of 0 for PSLMs. AUPRC is also more robust to imbalanced data than the traditional AUROC. Similar to AUROC, AUPRC ranges from 0 to 1, but unlike AUROC, AUPRC below 0.5 does not signify a worse-than-chance prediction: random guessing results in a data-dependent value above 0, which is taken into account and tested against.

Spearman’s *ρ* is regression statistic like Pearson’s r for measuring the strength of the relationship between continuous variables. Unlike Pearson’s r, which considers the linear relationship between variables, Spearman’s *ρ* only considers their rank-order, or monotonic correlation. In protein engineering applications, we are mostly concerned only with a model’s ability to rank mutations as being more or less stabilizing than one another, and since it does not involve information loss through binarization of labels or predictions, it may be informative than the previous statistics. *ρ* ranges from -1 for perfect reverse ordering, 0 for random ordering, and 1 for perfect ordering, according to:

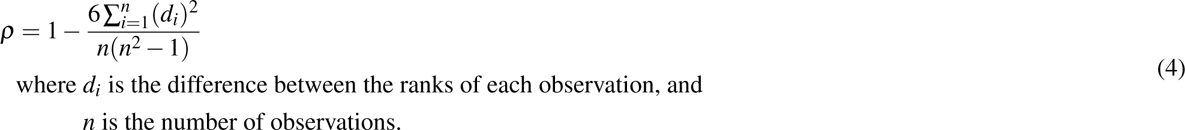

We also consider the mean and net stabilization, which collect all mutations whose predicted score (likelihood or stability) is greater than zero and simply take the mean or sum of the ground-truth label, respectively. The former statistic is related to the precision of the model, but also gives an estimate of the expected outcome of a positive prediction. Meanwhile, the latter statistic provides a good benchmark for model performance, since predictions must be both sensitive and precise, but the magnitude of the ground truth label is also important.

#### Per-Protein Aggregate Statistics

We suggest that, for protein engineering applications, the ability to correctly rank or classify mutations per-protein is more important than comparing the stability mutants of one protein relative to another. Therefore, we report per-protein statistics designated with the prefix “w” for (per-protein) weighted. In each case, we evaluate the statistic once for each set of mutations corresponding to a single wild-type structure, using all available mutations for that protein, and not counting proteins where the statistic is undefined. For instance, proteins with only one unique measurement value, even if it is shared by unique mutations, cannot be ranked, while proteins with no stabilizing mutations cannot be assessed using AUPRC. This statistical score is then weighted by the natural logarithm of the number of mutations contributing to that statistic, which factors in the reduced confidence in the significance for proteins with a small number of mutations. This decision carries the potential to overestimate the relative performance of methods trained only on larger collections of mutant stability measurements, such as Rosetta on parts of FireProtDB. All statistics are normalized by the weighted sum of all mutations considered such that the original domain of scores is retained, e.g. for weighted *ρ*:

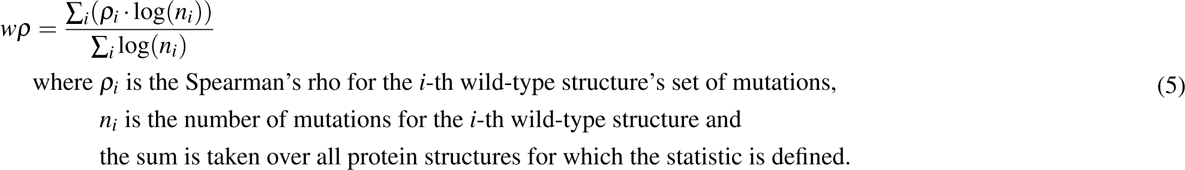

Finally, we consider the enrichment in higher-stability mutations when screening with stability models. We first consider the Normalized Discounted Cumulative Gain (NDCG) for this purpose. NDCG is a metric used to evaluate the effectiveness of information retrieval systems, especially in scenarios where top-ranked items are most important, such as in recommendation systems. This measure is derived from Discounted Cumulative Gain (DCG), which is the sum of the relevance scores (“gains”) of the retrieved items, discounted logarithmically for lower-ranked results. However, DCG is dependent on the particular set of results, so it is normalized against the ideal DCG (iDCG): a version of DCG where all the retrieved items are perfectly ranked by relevance. The ratio of DCG to iDCG is called NDCG. It ranges from 0 (worst) to 1 (best) and provides a relative measure of the performance of the ranking model, although there may be a limited distribution of NDCG scores, even for random predictions. In this context, we assign the relevance score to be the ground truth label, shifting the relevance by the value of the least stable protein, such that the most unstable protein has a relevance of 0:

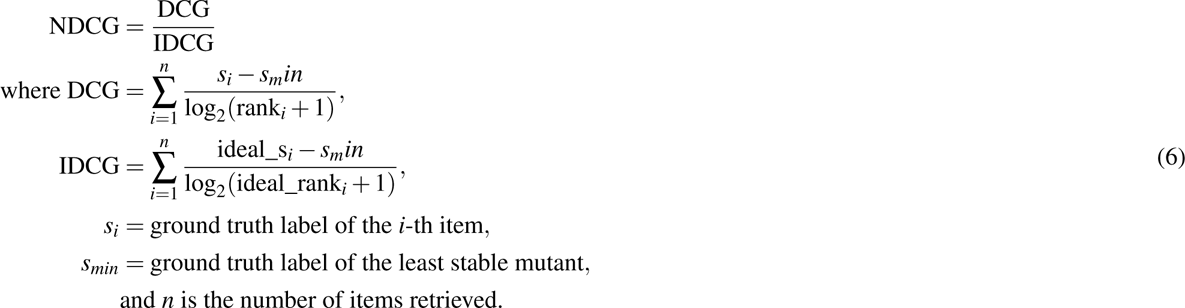

As such, the NDCG is like the Spearman *ρ* in that it depends on relative ranks, but the statistic depends more on the highest-stability mutations being among the top-ranked predictions (although the most stable mutations might not actually be stabilizing for every protein). Figure 6 below shows how the weights are assigned according to the predicted ranks with sets of different sizes, which is relevant to the per-protein weighted NDCG, which is the statistic we used as is calculated in the same way as for *ρ* in Eqn. 5. We do not consider NDCG for the whole dataset, as a small number of stabilizing mutations from only a few proteins contributes significantly to the score.

**Figure 6.**
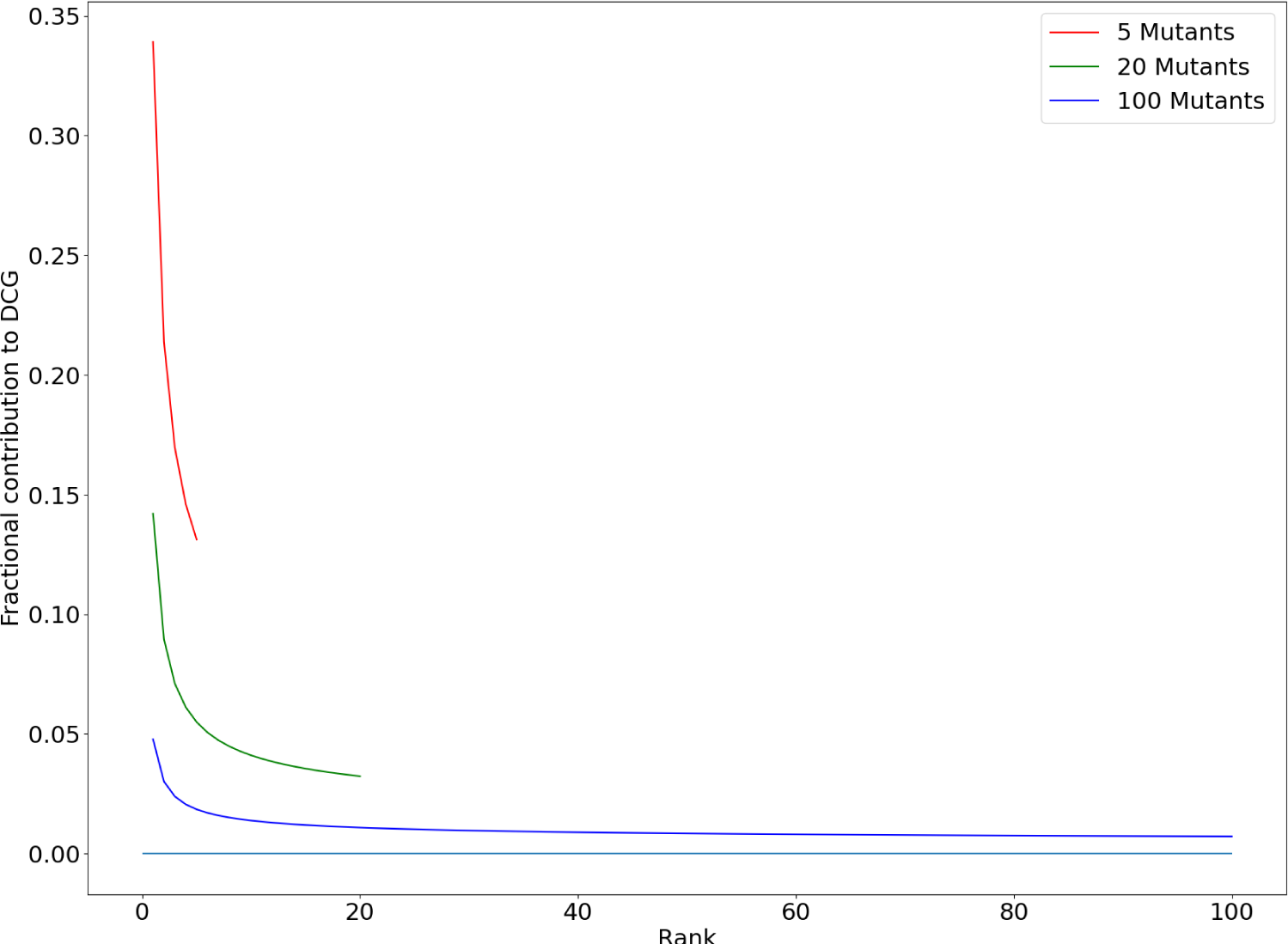
Scoring Contributions to NDCG by Predicted Rank. The weight associated with the relevance scores are shown for all predictions in hypothetical situations where a given protein has 5, 20, or 100 mutants, with the cyan line indicating zero.

We consider two additional strategies to quantify the detection of the most highly stabilizing mutations per-protein where the aggregation is performed differently. We first consider a percentile-based precision score, which assesses the fraction of each protein’s top-p percent most stabilizing predicted mutations which are experimentally stabilizing. p=(100-percentile)/100, such that the 95th percentile represents the top p=5% of predicted scores. The per-protein list of predictions is sorted, and the rounded-up percentage of the data length is used to get the top mutants from each sorted list. Rounding up is used to ensure each protein’s least-destabilizing variant is included when p is small. The number of mutations with labels above 0 are added to a running numerator per protein, while the length of the subset is added to a running denominator, which divides the numerator after all proteins are considered. We also consider a second strategy in which the recall of mutations is assessed per-protein when screening some multiple k of the number of experimentally stabilizing mutations for that protein. We evaluate multiples of k and report the fraction of all stabilizing mutations in the dataset identified when multiples of up to 3k top predictions are considered. The code to reproduce these statistics is available as per the Data Availability statement.

#### Bidirectional Statistics

For comparing predictions made for the same mutation in opposite directions (direct substitutions versus reversions to the wild-type from the mutant) we consider the antisymmetry and bias^55^. Antisymmetry is defined as the Pearson correlation coefficient between predictions of the direct and inverse mutations:

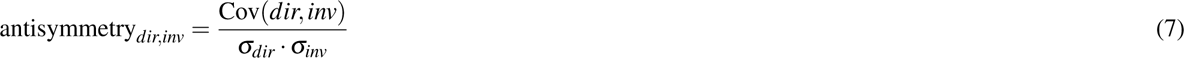

where *dir* and *inv* are the sets of direct and inverse mutation predictions, respectively. Since inverse mutations should always have equal magnitude and opposite sign to their direct counterparts, antisymmetry should be close to -1 for predictors which treat this physical reality correctly. In practice, achieving antisymmetric predictions involves either specialized training data (achieved through downsampling destabilizing mutations or augmenting datasets with hypothetical inverse mutations as in S669^55^, or using prediction strategies (usually based on the sequence) which are intrinsically antisymmetric. Similarly, bias quantifies the preference of models to designate mutations as destabilizing or stabilizing regardless of features:

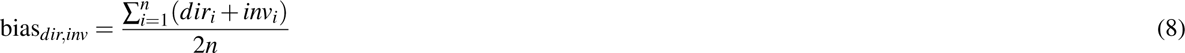

where *dir_i_* and *inv_i_* are the individual direct and inverse mutation predictions, respectively. Without taking precautions in training, most methods will reproduce the natural bias towards slight destabilization of mutants, which may be desirable, dependent on the context.

## Data Availability

Code for reproducing the experiments and analyses is available on Github: https://github.com/skalyaanamoorthy/thermostability-transfer. Tables with more extensive statistical analysis are available at the same location.

## Acknowledgements

The authors acknowledge the high-performance computing resources provided by the Digital Research Alliance of Canada. S.K. acknowledges the funding support from the NSERC Discovery Grant [RGPIN-2022-03348] and NFRF Exploration award [NFRFE-2019-01339]. S.R. acknowledges the NSERC CGS-M graduate scholarship.

## Author contributions statement

S.K. and S.R. conceived the project. S.R. designed and conducted the experiment(s), analyzed the results, and drafted the manuscript with input from S.K. S.K. supervised the project and acquired funding. All authors reviewed the manuscript.

