## Supplementary Table 1, Supplementary Table 2, Supplementary Table 3, Supplementary Table 4, Supplementary Table 5, Supplementary Figure 1-12 for "Zero-Shot Transfer of Protein Sequence Likelihood Models to Thermostability Prediction"

**Table 1.** Performance of Methods on FireProtDB with Neutral Mutations Removed

| | | | Whole-Dataset Statistics | | | Per-Protein Aggregate Statistics | | |
|---|---|---|---|---|---|---|---|---|
| measurement | type | model | MCC | AUPRC | $\rho$ | wNDCG | wAUPRC | w$\rho$ |
| $\Delta\Delta G$ | structural | Rosetta CartDDG | **0.454** | **0.523** | **0.549** | **0.903** | **0.732** | **0.369** |
| | structural | ESM-IF | 0.323 | 0.422 | 0.390 | 0.895 | 0.629 | 0.329 |
| | structural | KORPM | 0.356 | 0.411 | 0.457 | 0.891 | 0.658 | 0.317 |
| | structural | ESM-IF$_M$ | 0.306 | 0.414 | 0.384 | 0.892 | 0.606 | 0.316 |
| | structural | ProteinMPNN$_{20}$ | 0.376 | 0.427 | 0.400 | 0.899 | 0.702 | 0.300 |
| | structural | ProteinMPNN$_{30}$ | 0.365 | 0.425 | 0.391 | 0.896 | 0.704 | 0.295 |
| | structural | MIF-ST | 0.305 | 0.359 | 0.387 | 0.885 | 0.593 | 0.294 |
| | structural | MIF | 0.355 | 0.437 | 0.427 | 0.891 | 0.657 | 0.294 |
| | sequence(s) | MSA-T$_{mean}$ | 0.243 | 0.292 | 0.349 | 0.886 | 0.495 | 0.293 |
| | structural | ProteinMPNN$_{mean}$ | 0.401 | 0.441 | 0.403 | 0.898 | 0.712 | 0.289 |
| | sequence(s) | MSA-T$_1$ | 0.224 | 0.276 | 0.344 | 0.883 | 0.493 | 0.288 |
| | sequence(s) | Tranception | 0.197 | 0.219 | 0.232 | 0.882 | 0.510 | 0.277 |
| | structural | ProteinMPNN$_{10}$ | 0.374 | 0.420 | 0.384 | 0.891 | 0.684 | 0.261 |
| | sequence(s) | ESM-1V$_{mean}$ | 0.186 | 0.263 | 0.190 | 0.876 | 0.497 | 0.239 |
| | sequence(s) | ESM-1V$_2$ | 0.169 | 0.255 | 0.192 | 0.873 | 0.507 | 0.202 |
| | N/A | Gaussian noise | -0.001 | 0.135 | -0.020 | 0.822 | 0.414 | -0.052 |
| $\Delta T_m$ | structural | ESM-IF$_M$ | 0.398 | 0.650 | 0.636 | **0.902** | **0.692** | **0.367** |
| | structural | ESM-IF | 0.412 | **0.665** | **0.652** | 0.900 | 0.682 | 0.346 |
| | structural | Rosetta CartDDG | 0.397 | 0.574 | 0.602 | 0.899 | 0.734 | 0.328 |
| | structural | ProteinMPNN$_{30}$ | **0.459** | 0.636 | 0.514 | 0.893 | 0.710 | 0.285 |
| | structural | MIF-ST | 0.355 | 0.549 | 0.462 | 0.886 | 0.635 | 0.285 |
| | structural | KORPM | 0.332 | 0.459 | 0.448 | 0.882 | 0.648 | 0.279 |
| | structural | ProteinMPNN$_{20}$ | 0.457 | 0.650 | 0.512 | 0.895 | 0.707 | 0.276 |
| | structural | MIF | 0.384 | 0.592 | 0.509 | 0.886 | 0.641 | 0.275 |
| | structural | ProteinMPNN$_{mean}$ | 0.455 | 0.649 | 0.520 | 0.895 | 0.719 | 0.267 |
| | structural | ProteinMPNN$_{10}$ | 0.448 | 0.617 | 0.500 | 0.893 | 0.714 | 0.262 |
| | sequence(s) | MSA-T$_{mean}$ | 0.284 | 0.504 | 0.421 | 0.883 | 0.588 | 0.238 |
| | sequence(s) | MSA-T$_1$ | 0.289 | 0.511 | 0.417 | 0.882 | 0.590 | 0.209 |
| | sequence(s) | ESM-1V$_{mean}$ | 0.262 | 0.439 | 0.336 | 0.877 | 0.593 | 0.200 |
| | sequence(s) | Tranception | 0.269 | 0.468 | 0.431 | 0.873 | 0.613 | 0.183 |
| | sequence(s) | ESM-1V$_2$ | 0.244 | 0.444 | 0.355 | 0.873 | 0.590 | 0.179 |
| | N/A | Gaussian noise | -0.045 | 0.235 | 0.002 | 0.837 | 0.509 | 0.012 |

**Table 2.** Performance of Methods on FireProtDB with ProTherm Dataset Removed

| | | | Whole-Dataset Statistics | | | Per-Protein Aggregate Statistics | | |
|---|---|---|---|---|---|---|---|---|
| measurement | type | model | MCC | AUPRC | $\rho$ | wNDCG | wAUPRC | w$\rho$ |
| $\Delta\Delta G$ | structural | ProteinMPNN$_{30}$ | 0.183 | 0.542 | 0.384 | **0.917** | **0.741** | **0.449** |
| | structural | ProteinMPNN$_{20}$ | 0.192 | 0.548 | 0.384 | 0.916 | 0.729 | 0.446 |
| | structural | ProteinMPNN$_{mean}$ | 0.182 | 0.546 | 0.393 | 0.914 | 0.733 | 0.427 |
| | structural | ESM-IF | 0.157 | 0.497 | 0.335 | 0.904 | 0.698 | 0.387 |
| | structural | Rosetta CartDDG | **0.344** | **0.650** | **0.634** | 0.904 | 0.715 | 0.385 |
| | structural | MIF | 0.140 | 0.514 | 0.384 | 0.907 | 0.697 | 0.384 |
| | structural | ProteinMPNN$_{10}$ | 0.163 | 0.530 | 0.375 | 0.899 | 0.705 | 0.370 |
| | structural | ESM-IF$_M$ | 0.145 | 0.492 | 0.338 | 0.901 | 0.683 | 0.368 |
| | structural | KORPM | 0.321 | 0.593 | 0.520 | 0.885 | 0.669 | 0.333 |
| | structural | MIF-ST | 0.163 | 0.546 | 0.401 | 0.905 | 0.702 | 0.329 |
| | sequence(s) | MSA-T$_1$ | 0.056 | 0.448 | 0.220 | 0.894 | 0.637 | 0.305 |
| | sequence(s) | MSA-T$_{mean}$ | 0.077 | 0.459 | 0.225 | 0.896 | 0.638 | 0.304 |
| | sequence(s) | ESM-1V$_{mean}$ | 0.080 | 0.442 | 0.175 | 0.887 | 0.597 | 0.298 |
| | sequence(s) | ESM-1V$_2$ | 0.076 | 0.419 | 0.167 | 0.880 | 0.601 | 0.279 |
| | sequence(s) | Tranception | 0.028 | 0.378 | 0.114 | 0.88 | 0.613 | 0.223 |
| | N/A | Gaussian noise | 0.010 | 0.367 | 0.022 | 0.820 | 0.494 | -0.055 |
| $\Delta T_m$ | structural | MIF | 0.152 | 0.522 | 0.311 | **0.897** | 0.646 | **0.239** |
| | structural | ESM-IF$_M$ | 0.156 | 0.512 | 0.342 | 0.881 | 0.641 | **0.239** |
| | structural | ESM-IF | 0.152 | 0.514 | 0.354 | 0.885 | 0.644 | 0.219 |
| | structural | ProteinMPNN$_{mean}$ | 0.209 | 0.558 | **0.375** | 0.880 | 0.663 | 0.207 |
| | structural | ProteinMPNN$_{20}$ | 0.215 | **0.560** | 0.374 | 0.876 | 0.662 | 0.207 |
| | structural | ProteinMPNN$_{30}$ | 0.217 | 0.553 | 0.355 | 0.879 | 0.658 | 0.197 |
| | structural | ProteinMPNN$_{10}$ | **0.225** | 0.551 | 0.363 | 0.892 | **0.667** | 0.189 |
| | sequence(s) | ESM-1V$_{mean}$ | 0.107 | 0.470 | 0.198 | 0.869 | 0.636 | 0.153 |
| | structural | MIF-ST | 0.083 | 0.495 | 0.253 | 0.870 | 0.617 | 0.143 |
| | sequence(s) | MSA-T$_1$ | 0.087 | 0.480 | 0.205 | 0.870 | 0.626 | 0.136 |
| | structural | Rosetta CartDDG | 0.076 | 0.466 | 0.243 | 0.872 | 0.612 | 0.129 |
| | sequence(s) | Tranception | 0.084 | 0.473 | 0.257 | 0.866 | 0.631 | 0.127 |
| | sequence(s) | ESM-1V$_2$ | 0.091 | 0.463 | 0.195 | 0.869 | 0.634 | 0.116 |
| | sequence(s) | MSA-T$_{mean}$ | 0.093 | 0.472 | 0.208 | 0.861 | 0.619 | 0.111 |
| | structural | KORPM | 0.111 | 0.441 | 0.208 | 0.861 | 0.609 | 0.089 |
| | N/A | Gaussian noise | -0.067 | 0.389 | -0.054 | 0.839 | 0.579 | -0.061 |

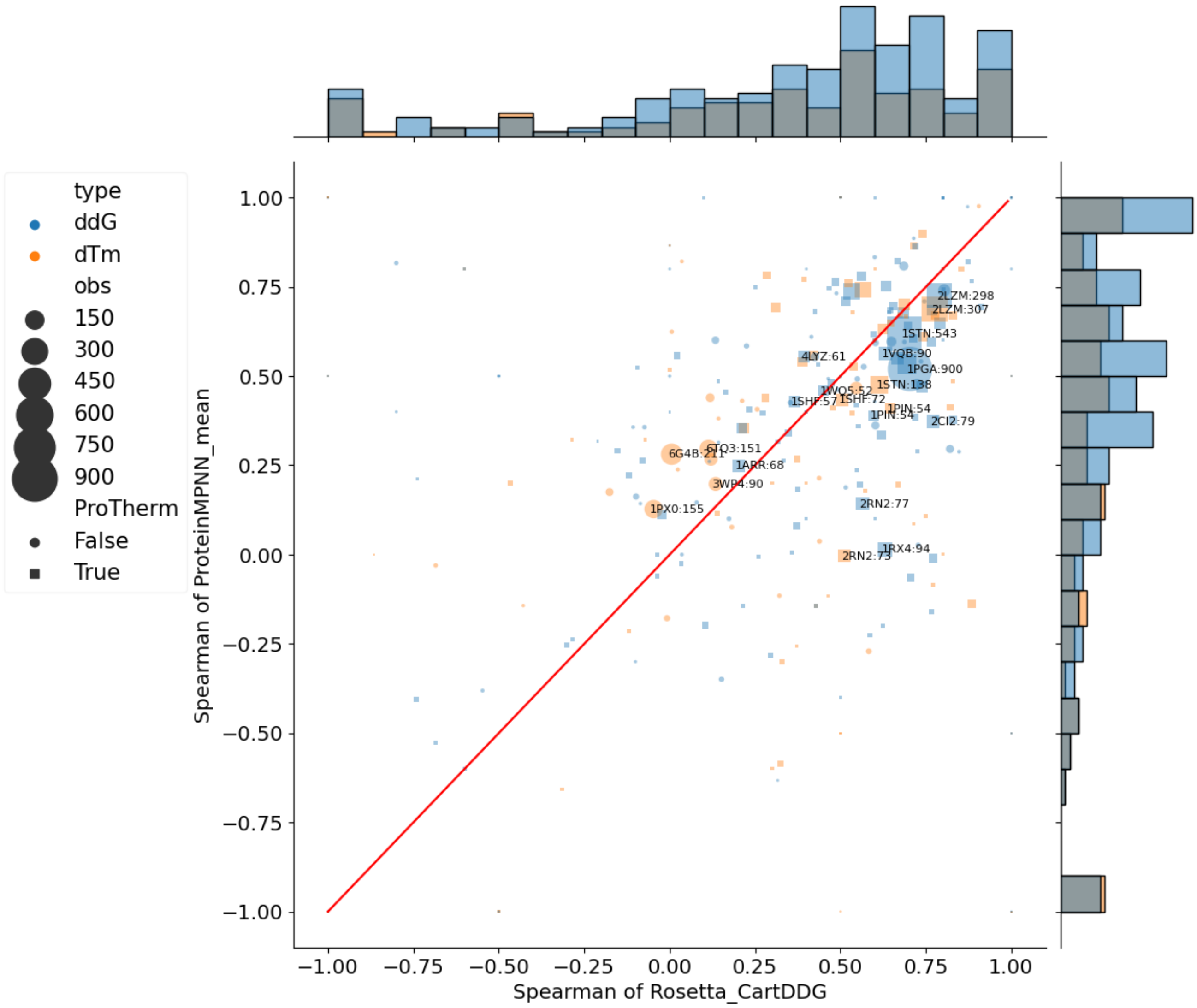


**Figure 1. Per-Protein Comparison of Ranking Performance of Top Models**. Each point on the plot indicates the Spearman $\rho$ of each of the corresponding models on the axes. The red diagonal line helps to visualize improved performance: points above the diagonal indicate that ProteinMPNN had improved performance. The size of each point is related to the number of mutations for that wild-type structure, which were used to measure $\rho$. Square point indicate proteins not found in the ProTherm dataset. Some points are labelled with the PDB code of the structure used and the number of mutations included. The marginals indicate the unweighted sum of points in their corresponding binned row / column for the model on the same axis.

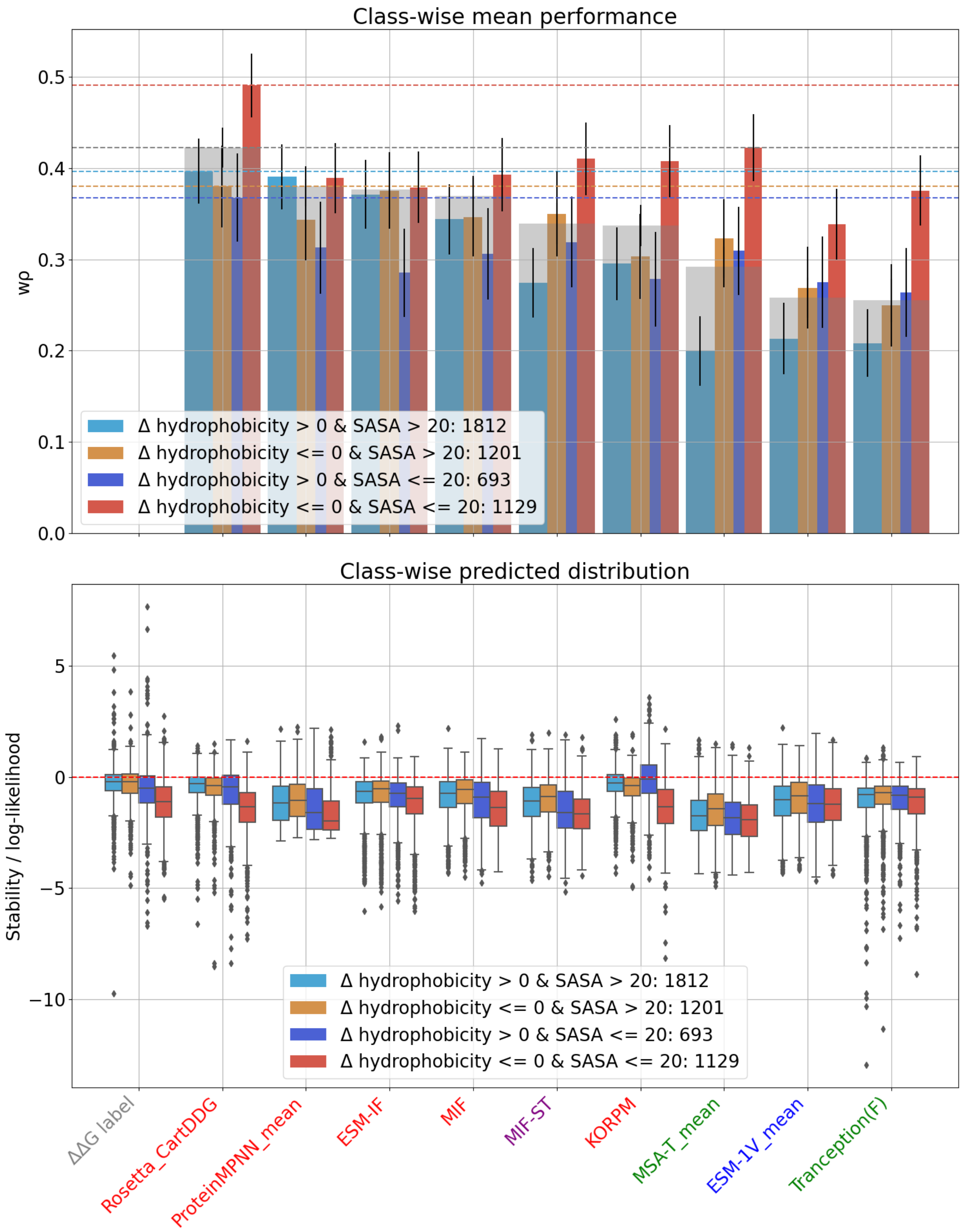


**Figure 2. Per-Protein Ranking Performance of Different Physicochemical Mutation Classes**. The FireProtDB unique mutations are divided into four groups with depending on their change in hydrophobicity and the wild-type solvent accessible surface area (SASA) while the grey overlaid bar indicates the non-split performance. The top plot shows the distribution of weighted Spearman $\rho$ across 1000 bootstrapped replicates (error bar) while the bottom shows the predicted stability for each class. SASA in Å$^2$ is calculated by DSSP[1], while change in hydrophobicity is based on the Kyte-Doolittle hydrophobicity scale[2], with a positive value indicating the hydrophobicity has increased. The bottom plot shows the distribution over all predictions, with the ground truth label distribution at the left. The red horizontal dashed line indicates the decision threshold used for binary classification with MCC.

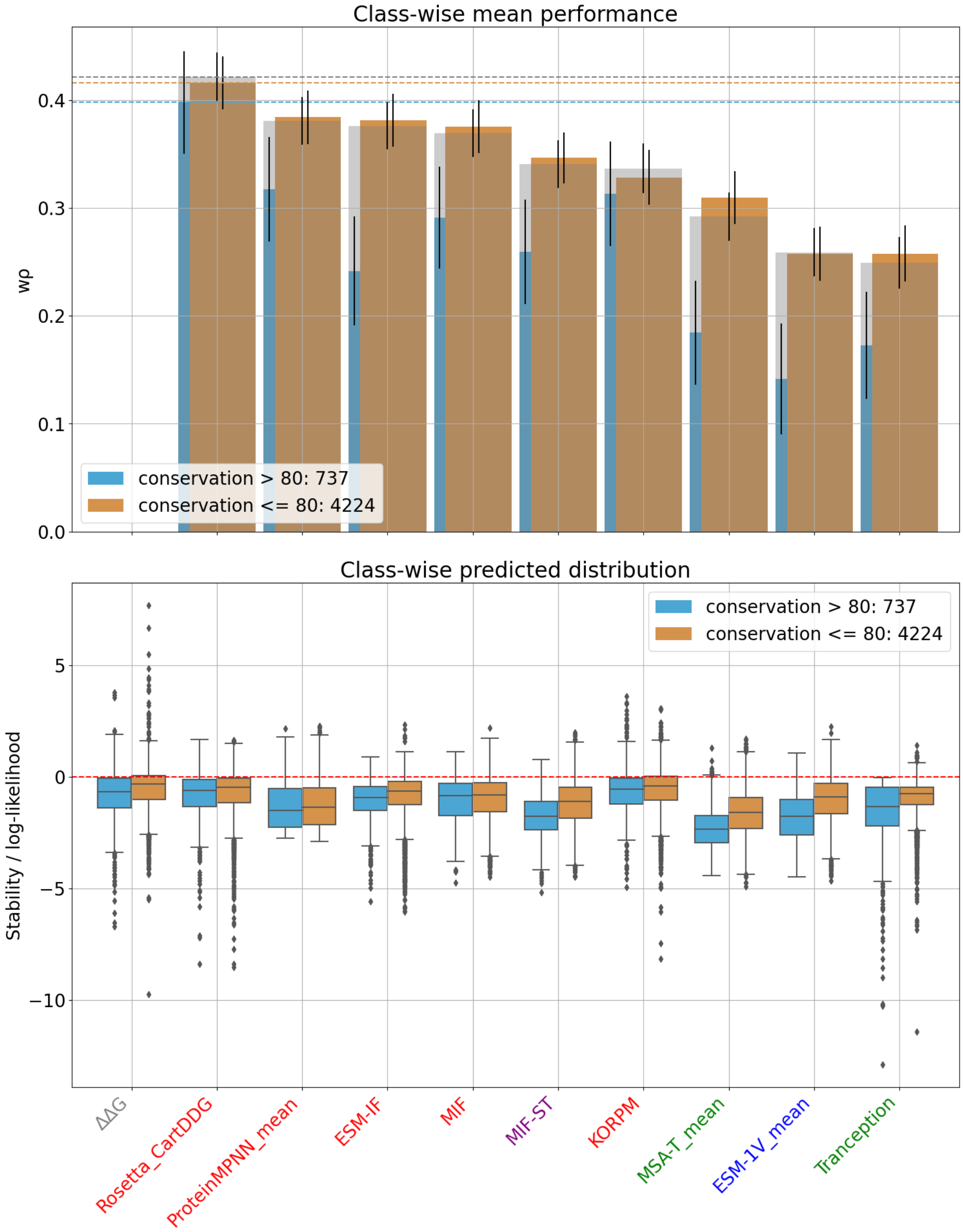


**Figure 3. Per-Protein Ranking Performance with Different Aligned Position Conservation**. The FireProtDB unique mutations are divided into groups based on the percent conservation of the wild-type residue at the mutated position of the alignment used by evolutionary methods, while the grey overlaid bar indicates the non-split performance. The top plot shows the distribution of weighted Spearman $\rho$ across 1000 bootstrapped replicates (error bar) while the bottom shows the predicted stability for each class.

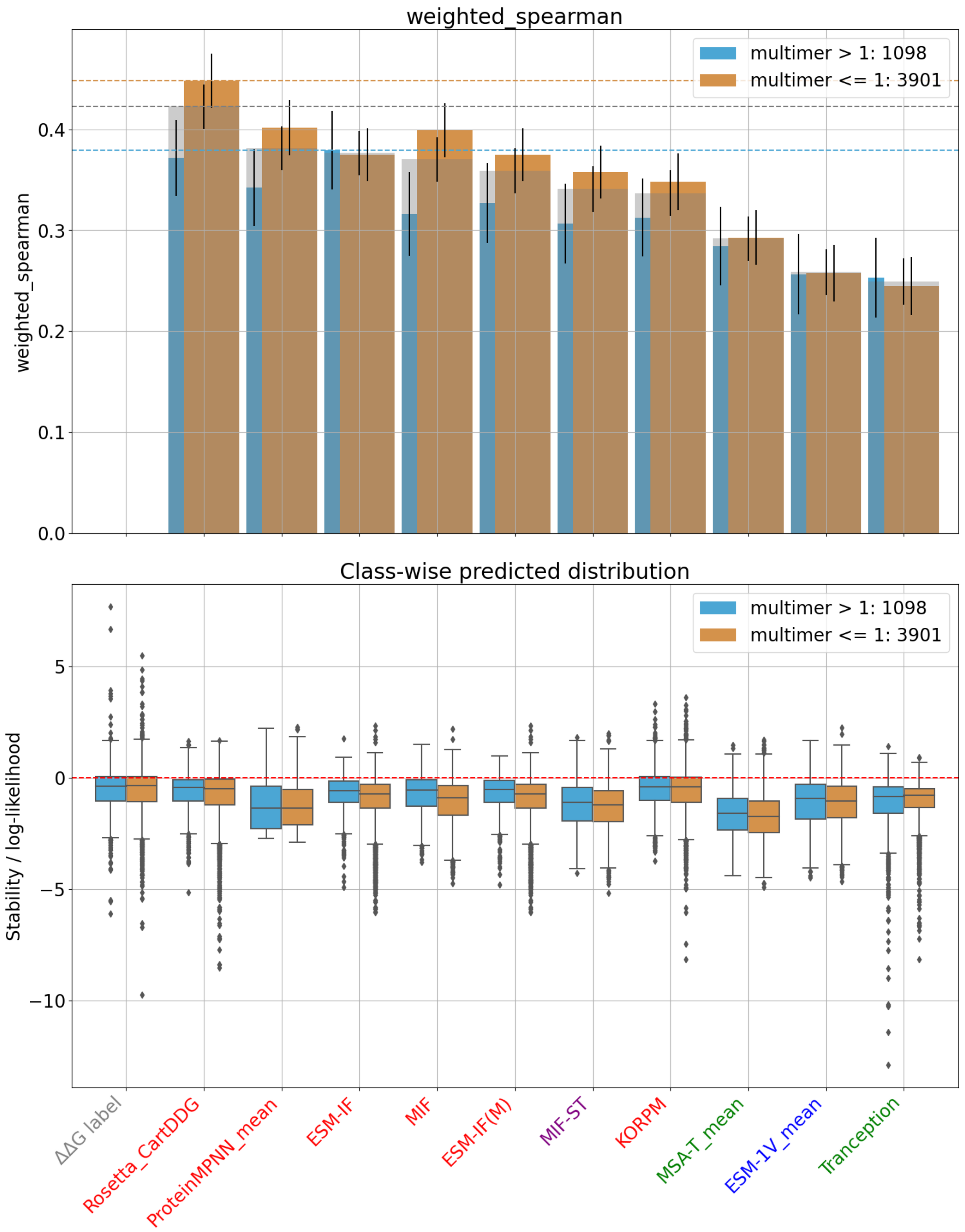


**Figure 4. Per-Protein Ranking Performance with Quaternary Structure**. The FireProtDB unique mutations are divided on the basis of quaternary structure: whether the given protein structure is monomeric or multimeric, while the grey overlaid bar indicates the non-split performance. The top plot shows the distribution of weighted Spearman $\rho$ across 1000 bootstrapped replicates (error bar) while the bottom shows the predicted stability for each class.

**Table 3.** Stability Engineering Statistics Performance of Methods on FireProtDB. TP: true postives; FP: false positives; TN: true negatives; FN: false negatives, AUPPC: area under the percentile-wise precision curve; AUKXRC: area under the K-times recovery curve; Mean T1S: mean stability of each protein's top-scoring mutant; Mean Stab.: mean stability of all predicted-stabilizing mutations; Net Stab.: net stabilizing effect of all predicted stabilizing mutations, combined.

| meas | model | TP | FP | TN | FN | AUPPC | AUKXRC | Mean T1S | Mean Stab. | Net Stab. |
|---|---|---|---|---|---|---|---|---|---|---|
| $\Delta\Delta G$ | $\Delta\Delta G$ | 1370 | 0 | 3672 | 0 | 0.562 | 0.495 | 1.069 | 0.771 | 1056.73 |
| | $\Delta T_m$ | 376 | 73 | 1053 | 52 | 0.531 | 0.451 | 0.732 | 0.624 | 280.11 |
| | ProteinMPNN$_{mean}$ | 370 | 263 | 3409 | 1000 | **0.424** | **0.385** | 0.033 | 0.144 | **91.375** |
| | Rosetta_CartDDG | 577 | 457 | 3215 | 793 | 0.417 | 0.383 | -0.033 | 0.072 | 74.53 |
| | ProteinMPNN$_{10}$ | 342 | 264 | 3408 | 1028 | 0.415 | 0.378 | -0.021 | 0.094 | 57.175 |
| | ProteinMPNN$_{20}$ | 387 | 300 | 3372 | 983 | 0.421 | 0.382 | 0.06 | 0.082 | 56.04 |
| | MIF-ST | 239 | 190 | 3482 | 1131 | 0.383 | 0.355 | -0.018 | 0.122 | 52.54 |
| | MSA-T$_{mean}$ | 128 | **85** | **3587** | 1242 | 0.349 | 0.329 | -0.081 | **0.172** | 36.665 |
| | MSA-T$_1$ | 134 | 113 | 3559 | 1236 | 0.346 | 0.327 | -0.127 | 0.069 | 17.16 |
| | ProteinMPNN$_{30}$ | 390 | 312 | 3360 | 980 | 0.417 | 0.379 | -0.046 | 0.021 | 14.92 |
| | Tranception | 86 | **85** | **3587** | 1284 | 0.333 | 0.318 | -0.256 | 0.081 | 13.925 |
| | ESM-IF | 284 | 228 | 3444 | 1086 | 0.404 | 0.371 | 0.019 | 0.021 | 10.645 |
| | MIF | 353 | 348 | 3324 | 1017 | 0.403 | 0.369 | **0.074** | -0.009 | -6.365 |
| | ESM-IF$_M$ | 282 | 240 | 3432 | 1088 | 0.402 | 0.37 | -0.054 | -0.024 | -12.775 |
| | ESM-1V$_{mean}$ | 228 | 330 | 3342 | 1142 | 0.342 | 0.325 | -0.279 | -0.329 | -183.755 |
| | KORPM | **629** | 685 | 2987 | **741** | 0.387 | 0.36 | -0.126 | -0.141 | -185.11 |
| | ESM-1V$_2$ | 224 | 329 | 3343 | 1146 | 0.338 | 0.323 | -0.29 | -0.376 | -208.17 |
| | Gaussian noise | 689 | 1885 | 1787 | 681 | 0.262 | 0.273 | -0.734 | -1.046 | -2693.42 |
| $\Delta T_m$ | $\Delta T_m$ | 908 | 0 | 1880 | 0 | 0.622 | 0.491 | 4.354 | 3.148 | 2858.025 |
| | $\Delta\Delta G$ | 376 | 52 | 1053 | 73 | 0.554 | 0.464 | 4.116 | 2.888 | 1235.85 |
| | ProteinMPNN$_{mean}$ | 339 | 209 | 1671 | 569 | **0.444** | **0.372** | 1.155 | 1.44 | **789.11** |
| | ProteinMPNN$_{30}$ | 370 | 240 | 1640 | 538 | 0.443 | 0.371 | 0.963 | 1.221 | 744.695 |
| | ProteinMPNN$_{20}$ | 353 | 225 | 1655 | 555 | 0.442 | **0.372** | 1.076 | 1.284 | 742.425 |
| | ProteinMPNN$_{10}$ | 323 | 203 | 1677 | 585 | 0.438 | 0.369 | 1.024 | 1.315 | 691.605 |
| | ESM-IF | 238 | 154 | 1726 | 670 | 0.419 | 0.359 | **1.378** | **1.652** | 647.47 |
| | ESM-IF$_M$ | 244 | 155 | 1725 | 665 | 0.419 | 0.358 | 1.075 | 1.49 | 594.51 |
| | MIF-ST | 238 | 193 | 1687 | 670 | 0.404 | 0.351 | 0.668 | 0.839 | 361.695 |
| | MIF | 317 | 245 | 1635 | 591 | 0.42 | 0.36 | 1.048 | 0.643 | 361.535 |
| | Rosetta_CartDDG | 441 | 447 | 1433 | 467 | 0.408 | 0.353 | 0.607 | 0.384 | 341.185 |
| | MSA-T$_1$ | 194 | 154 | 1726 | 714 | 0.389 | 0.339 | 0.403 | 0.652 | 226.82 |
| | MSA-T$_{mean}$ | 193 | **153** | **1727** | 715 | 0.392 | 0.341 | 0.543 | 0.655 | 226.69 |
| | Tranception | 179 | 173 | 1707 | 729 | 0.373 | 0.331 | -0.138 | 0.163 | 57.205 |
| | ESM-1V$_{mean}$ | 265 | 280 | 1600 | 643 | 0.384 | 0.339 | 0.027 | -0.09 | -48.97 |
| | ESM-1V$_2$ | 252 | 283 | 1597 | 656 | 0.379 | 0.337 | -0.324 | -0.213 | -113.8 |
| | KORPM | **448** | 477 | 1403 | **460** | 0.4 | 0.35 | 0.192 | -0.177 | -163.73 |
| | Gaussian noise | 470 | 939 | 941 | 438 | 0.32 | 0.302 | -1.614 | -2.866 | -4037.73 |

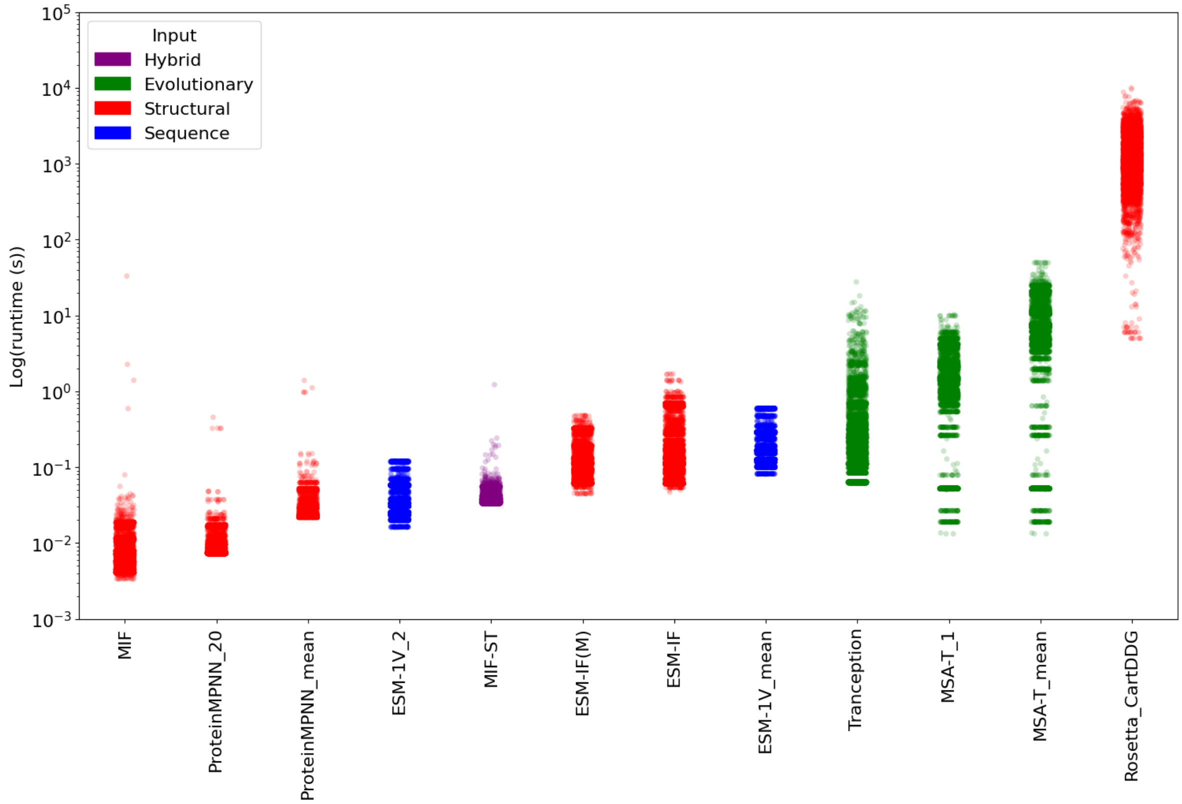


**Figure 5. Individual Mutant Prediction Times for FireProtDB.** Each point represents the time taken in log(seconds) to make a stability prediction for a single mutation; Tranception's time are estimated by dviding the total time per protein by the number of mutants. Points are colored according to the method's type. KORPM has an estimated constant runtime of 1 millisecond per mutant.

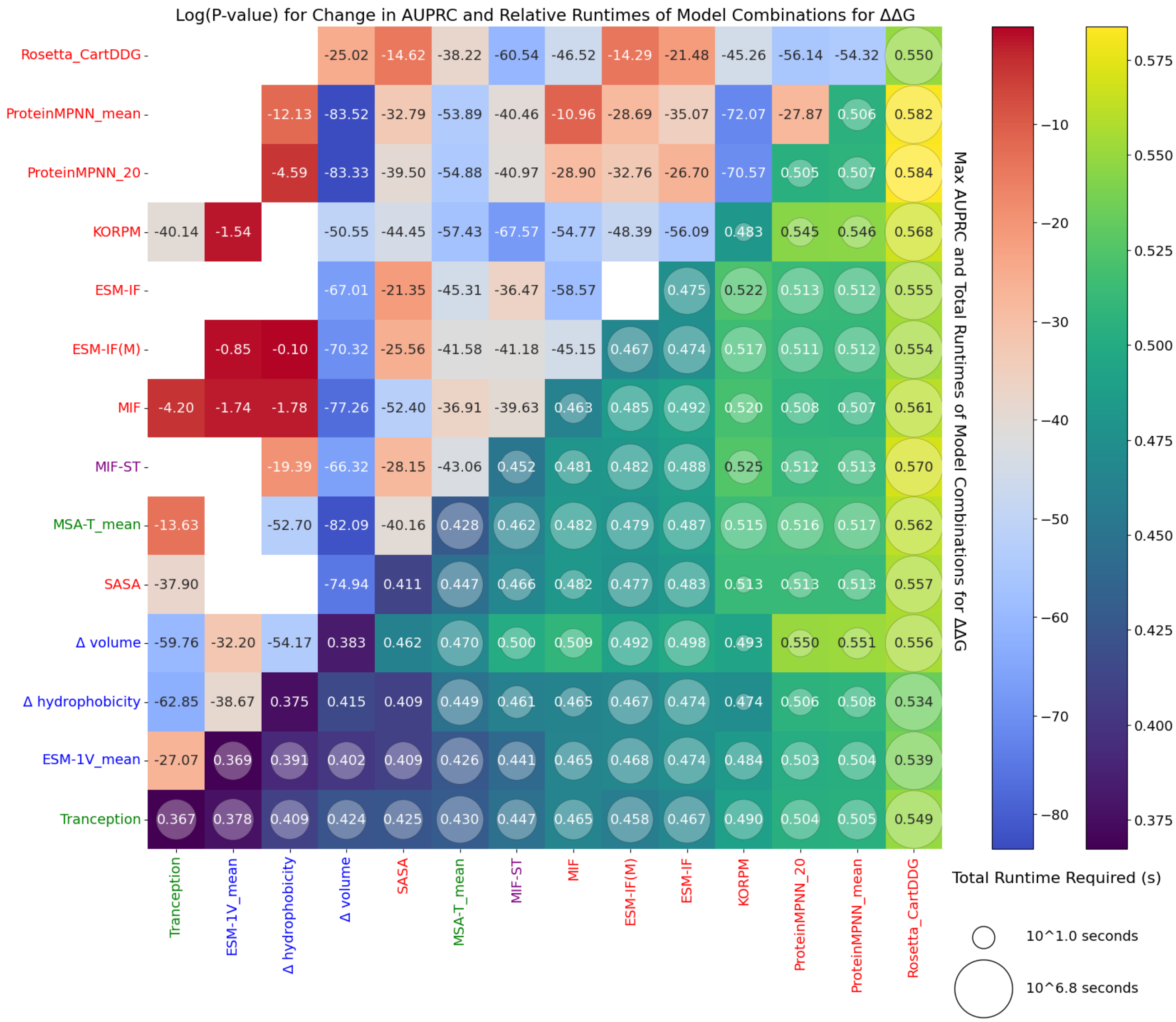


**Figure 6. Whole-Dataset Classification Performance and Runtimes of 2-Model Combinations on FireProtDB**. The annotations and colors in the upper left of the plot indicate the log(P-value) for improvement in AUPRC (ungrouped) for the best of five weighted combinations of normalized predictions of two models, compared to the best constituent model (100 bootstrapped replicates). White cells indicate that performance of all combinations was reduced. Performance of individual models is given on the diagonal. On the lower triangle, the size of the circular marker's radius is proportional to the log(runtime) for making both sets of predictions. The color and cell annotation indicate the AUPRC for the best model combinations.

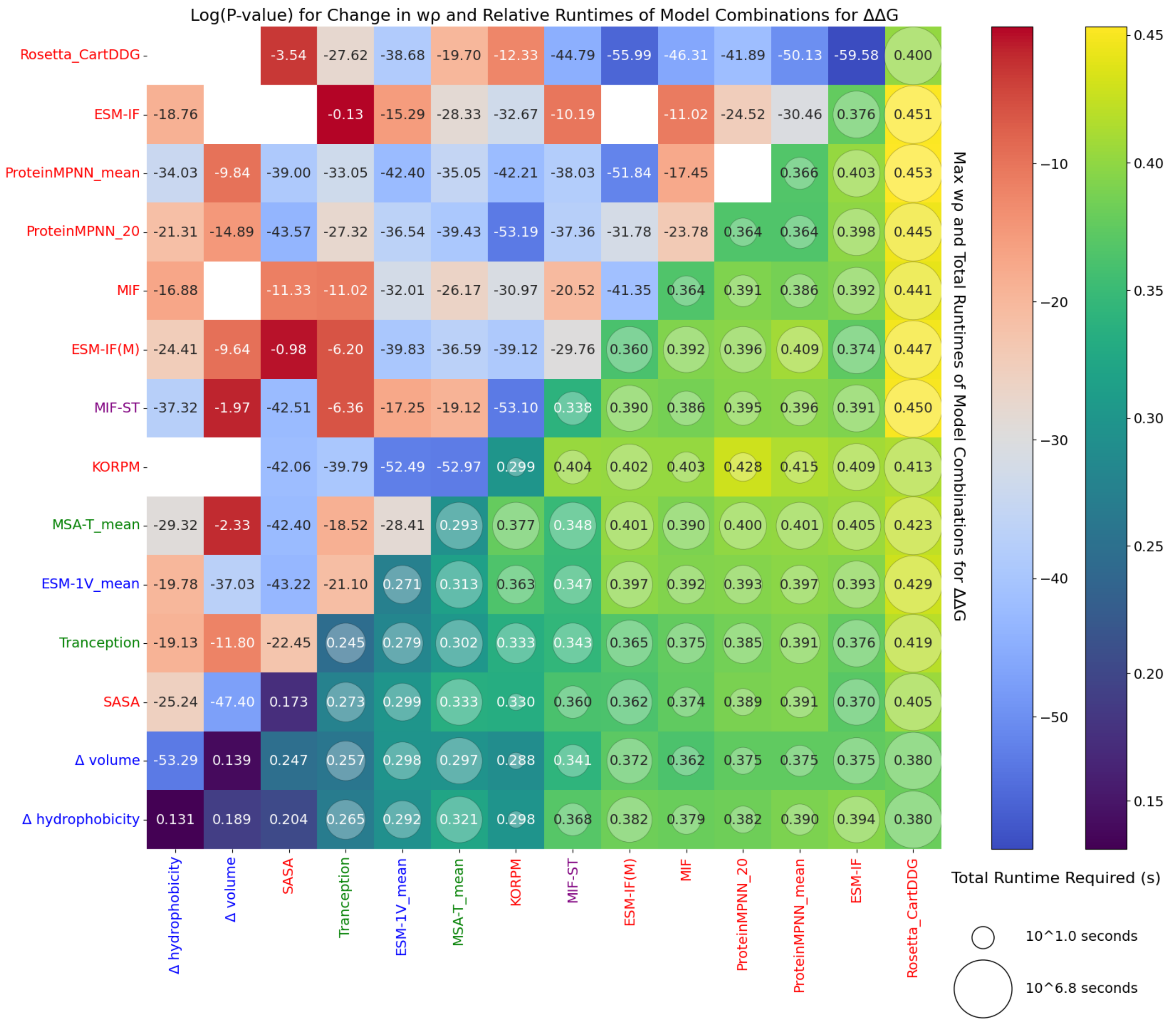


**Figure 7. Per-Protein Ranking Performance and Runtimes of 2-Model Combinations on FireProtDB**. The annotations and colors in the upper left of the plot indicate the log(P-value) for improvement in weighted Spearman $\rho$ for the best of five weighted combinations of normalized predictions of two models, compared to the best constituent model (100 bootstrapped replicates). White cells indicate that performance of all combinations was reduced. Average performance of individual models is given on the diagonal. On the lower triangle, the size of the circular marker's radius is proportional to the log(runtime) for making both sets of predictions. The color and cell annotation indicate the weighted Spearman $\rho$ for the best model combinations. X-axis labels are color-coded as grey: label, red: structural, blue: sequence-only, purple: structural and sequence features, green: evolutionary.

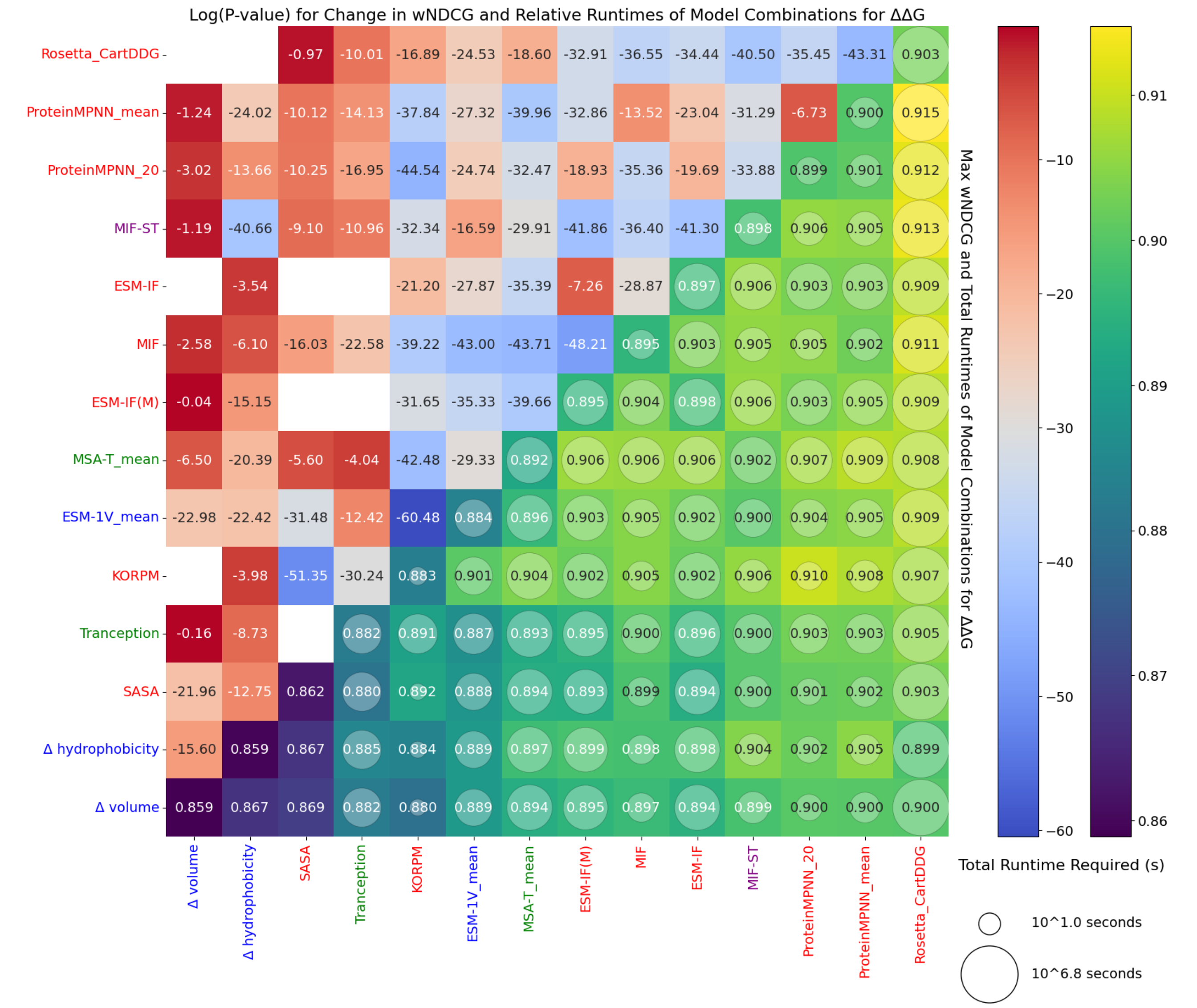


**Figure 8. Per-Protein Screening Prioritization Performance and Runtimes of 2-Model Combinations on FireProtDB.** The annotations and colors in the upper left of the plot indicate the log(P-value) for improvement in weighted NDCG for the best of five weighted combinations of normalized predictions of two models, compared to the best constituent model (100 bootstrapped replicates). White cells indicate that performance of all combinations was reduced. Average performance of individual models is given on the diagonal. On the lower triangle, the size of the circular marker's radius is proportional to the log(runtime) for making both sets of predictions. The color and cell annotation indicate the weighted NDCG for the best model combinations. X-axis labels are color-coded as grey: label, red: structural, blue: sequence-only, purple: structural and sequence features, green: evolutionary.

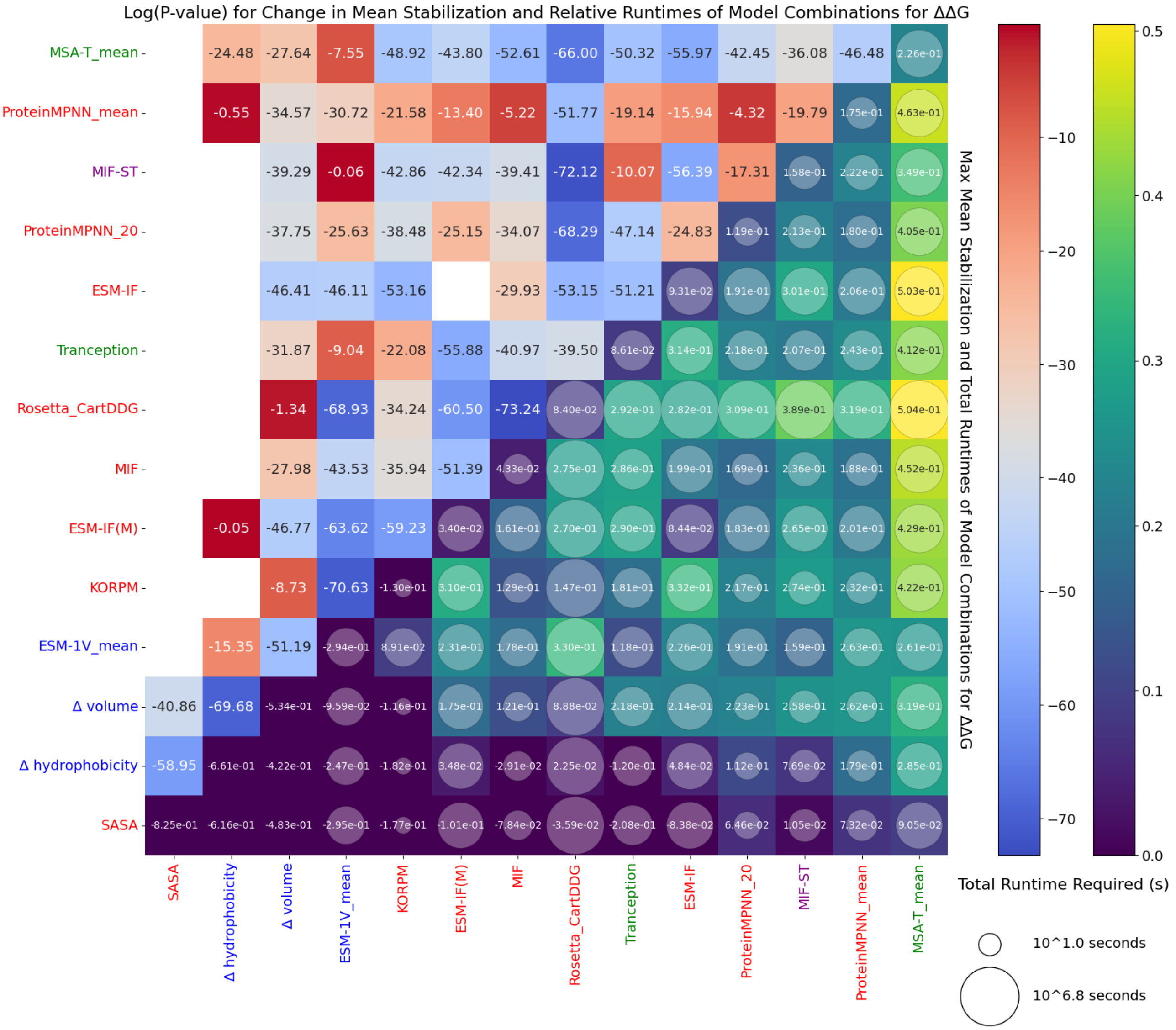


**Figure 9.** **Average Stabilization and Runtimes of 2-Model Combinations on FireProtDB**. The annotations and colors in the upper left of the plot indicate the log(P-value) for improvement in mean stabilization per screened mutation (kcal/mol) for the best of five weighted combinations of normalized predictions of two models, compared to the best constituent model (paired t-test, 100 bootstrapped replicates). White cells indicate that performance of all combinations was reduced. Average performance of individual models is given on the diagonal. On the lower triangle, the size of the circular marker's radius is proportional to the log(runtime) for making both sets of predictions. The color and cell annotation indicate the performance for the best model combinations. X-axis labels are color-coded as grey: label, red: structural, blue: sequence-only, purple: structural and sequence features, green: evolutionary.

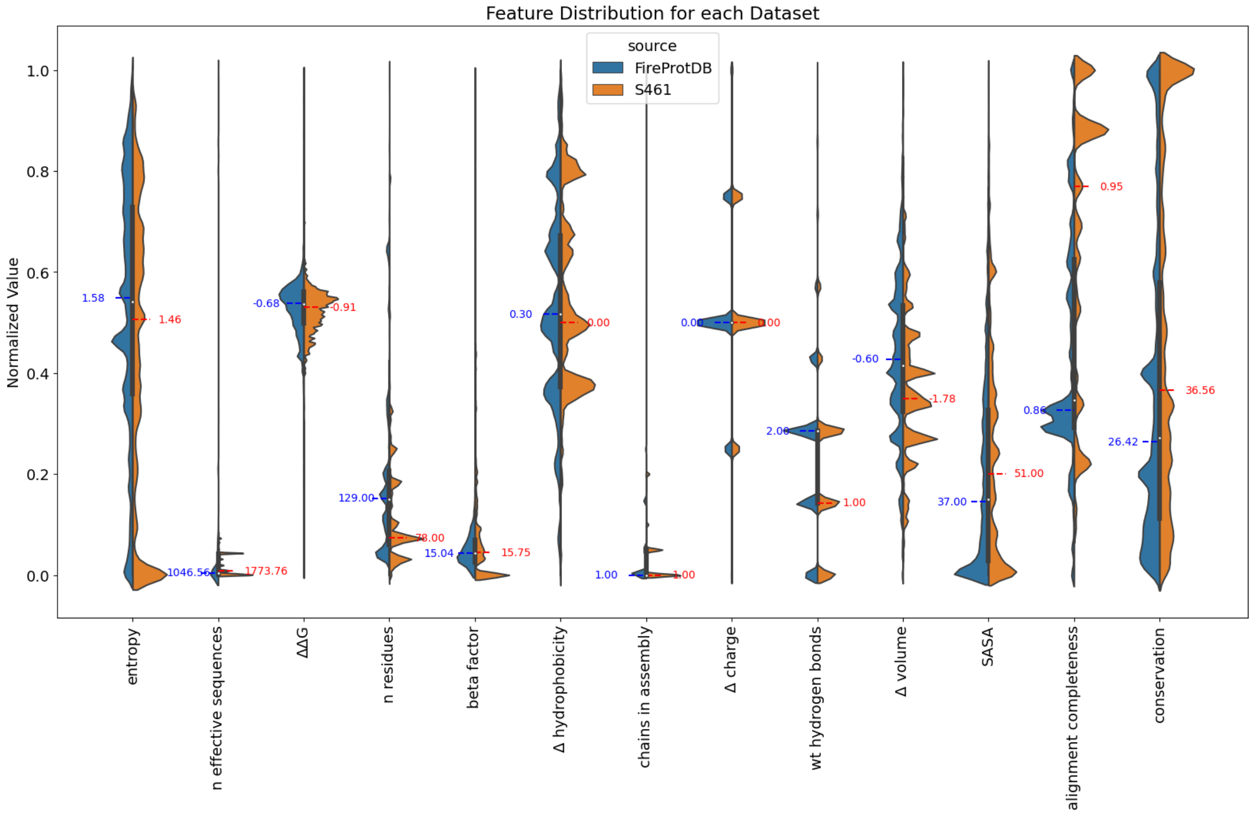


**Figure 10. Feature Distributions for Tested Datasets**. Each violin represents an extracted feature across all unique mutations in the FireProtDB (blue) or S461 (orange) datasets. The features have been rescaled from 0 to 1 for comparison, but the median values of their unnormalized distributions are given in colors corresponding to the distribution. Violin smoothing inaccurately reflects discrete values.

**Table 4.** Selection of Top Performing Pairs Across Stability Engineering Statistics for Direct Mutations on S461

| Model 1 | Model 2 | Weight 2 | wNDCG | AUPPC | Net Stabilization |
|---|---|---|---|---|---|
| $\Delta\Delta G$ | $\Delta\Delta G$ | 0 | 1 | 0.416336 | 36.998 |
| Rosetta_CartDDG | ESM-1V$_{mean}$ | 1 | **0.931028** | 0.296098 | 7.953 |
| ESM-1V$_{mean}$ | KORPM | 1 | 0.930481 | 0.286265 | 7.056 |
| KORPM | MIF-ST | 0.5 | 0.92831 | 0.285743 | -1.8 |
| ACDC-NN | ESM-1V$_{mean}$ | 0.5 | 0.927316 | 0.2754 | -1.681 |
| MAESTRO | ESM-1V$_{mean}$ | 1 | 0.926112 | 0.280795 | 7.712 |
| KORPM | Tranception | 1 | 0.925958 | 0.270542 | **14.164** |
| ESM-1V$_{mean}$ | PopMusic | 0.2 | 0.925704 | 0.291407 | 2.47 |
| Tranception | Rosetta_CartDDG | 0.5 | 0.92551 | 0.283309 | 9.924 |
| ESM-1V$_{mean}$ | SASA | 0.2 | 0.924505 | 0.270943 | -30.6 |
| ESM-IF$_M$ | ACDC-NN | 0.5 | 0.923867 | 0.281231 | -1.102 |
| Rosetta_CartDDG | PremPS | 0.5 | 0.919268 | **0.310211** | 8.007 |
| DDGun3D | MIF-ST | 1 | 0.907524 | 0.308517 | 6.607 |
| Rosetta_CartDDG | ProteinMPNN$_{20}$ | 0.5 | 0.920423 | 0.305529 | 2.873 |
| Rosetta_CartDDG | DDGun3D | 0.2 | 0.913293 | 0.304084 | -0.815 |
| MIF | DDGun3D | 0.5 | 0.895255 | 0.304036 | 6.652 |
| Rosetta_CartDDG | PopMusic | 0.2 | 0.918531 | 0.303533 | 6.711 |
| Rosetta_CartDDG | ProteinMPNN$_{mean}$ | 0.5 | 0.916222 | 0.303288 | 5.793 |
| DDGun | MIF | 1 | 0.894002 | 0.302926 | 5.31 |
| DDGun | MIF-ST | 0.5 | 0.906197 | 0.301848 | 7.221 |
| DDGun | DUET | 0.5 | 0.883613 | 0.301511 | -7.99 |
| PremPS | ESM-IF$_M$ | 1 | 0.917239 | 0.27653 | 14.106 |
| ESM-IF | PremPS | 0.5 | 0.904751 | 0.266945 | 13.046 |
| MIF-ST | PremPS | 0.5 | 0.90437 | 0.284387 | 12.242 |
| Rosetta_CartDDG | ESM-IF$_M$ | 1 | 0.922209 | 0.29855 | 11.774 |
| INPS3D | MIF-ST | 1 | 0.911253 | 0.289014 | 11.08 |
| PremPS | MIF | 0.2 | 0.903017 | 0.287048 | 10.575 |
| ESM-1V$_{mean}$ | PremPS | 0.2 | 0.910585 | 0.269041 | 10.446 |
| ESM-1V$_{mean}$ | MIF | 0.5 | 0.91697 | 0.294287 | 10.429 |
| Rosetta_CartDDG | ESM-IF | 1 | 0.918898 | 0.296712 | 10.274 |

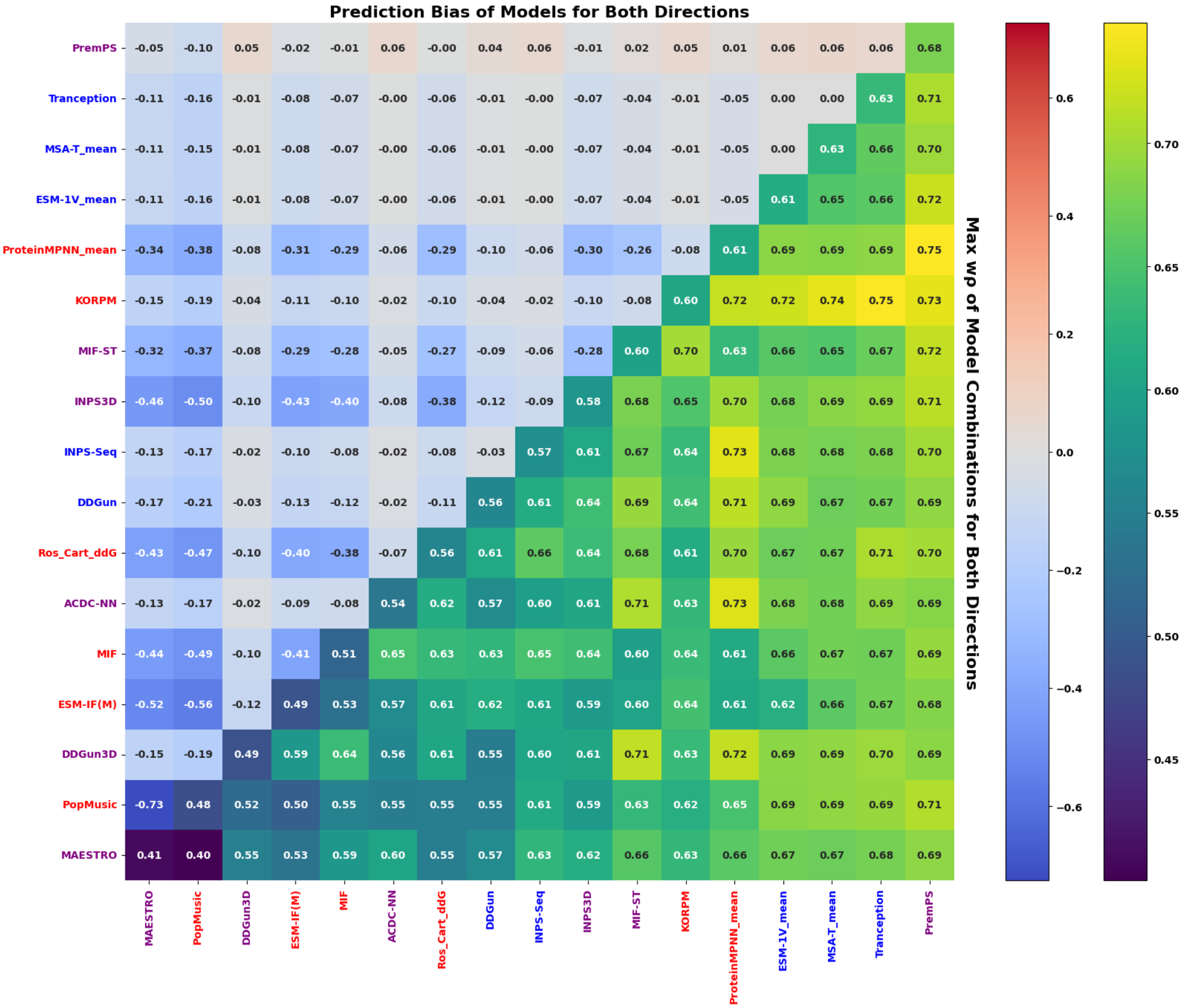


**Figure 11.** **Per-Protein Ranking Performance and Bias of 2-Model Combinations on S461 (Both Directions)**. The annotations and colors in the upper left of the plot indicate the bias between matched direct and reversion mutant predictions of high-performing models. Performance of individual models in terms of weighted Spearman $\rho$ (w$\rho$) is given on the diagonal, while the lower triangle indicates w$\rho$ for the best of five weighted combinations of normalized predictions of two models

**Table 5.** Selection of Top Performing Pairs Across Stability Engineering Statistics for Bidirectional Mutations on S461

| Model 1 | Model 2 | Weight 2 | wNDCG | AUPPC | Net Stabilization |
|---|---|---|---|---|---|
| $\Delta\Delta G$ | $\Delta\Delta G$ | 0 | 1 | 0.853722 | 601.828 |
| KORPM | $\text{ProteinMPNN}_{20}$ | 0.5 | **0.953748** | 0.787994 | 497.362 |
| PremPS | $\text{ProteinMPNN}_{mean}$ | 1 | 0.953738 | 0.805787 | 533.997 |
| PremPS | $\text{ProteinMPNN}_{20}$ | 1 | 0.953735 | **0.806188** | 535.622 |
| KORPM | $\text{ProteinMPNN}_{mean}$ | 0.5 | 0.952828 | 0.789697 | 494.41 |
| KORPM | $\text{MSA-T}_{mean}$ | 0.5 | 0.952308 | 0.795426 | 537.93 |
| Tranception | KORPM | 0.5 | 0.94986 | 0.796212 | **544.317** |
| $\text{ESM-1V}_{mean}$ | KORPM | 1 | 0.949717 | 0.797905 | 541.193 |
| PremPS | KORPM | 1 | 0.949269 | 0.801664 | 526.134 |
| KORPM | MIF-ST | 1 | 0.949183 | 0.78771 | 499.533 |
| DDGun3D | $\text{ProteinMPNN}_{mean}$ | 1 | 0.948286 | 0.804225 | 522.756 |
| DDGun | $\text{ProteinMPNN}_{mean}$ | 1 | 0.945023 | 0.805947 | 507.34 |
| $\text{ProteinMPNN}_{20}$ | DDGun | 0.5 | 0.943463 | 0.803185 | 504.723 |
| PremPS | MIF | 0.2 | 0.942322 | 0.80107 | 529.937 |
| ACDC-NN | $\text{ProteinMPNN}_{mean}$ | 1 | 0.947969 | 0.800984 | 533.751 |
| DDGun3D | MIF-ST | 1 | 0.948267 | 0.800866 | 526.408 |
| $\text{ProteinMPNN}_{20}$ | ACDC-NN | 0.5 | 0.946904 | 0.800785 | 529.062 |
| PremPS | $\text{ESM-1V}_{mean}$ | 1 | 0.946865 | 0.797348 | 542.249 |
| Tranception | ThermoNet | 0.5 | 0.935456 | 0.781761 | 538.12 |
| $\text{ESM-1V}_{mean}$ | Tranception | 0.2 | 0.935601 | 0.770283 | 537.512 |
| Tranception | Tranception | 0 | 0.92492 | 0.759358 | 537.152 |
| ACDC-NN | MIF-ST | 1 | 0.946382 | 0.797738 | 535.644 |
| $\text{ESM-1V}_{mean}$ | $\text{ProteinMPNN}_{20}$ | 1 | 0.942561 | 0.78875 | 535.395 |

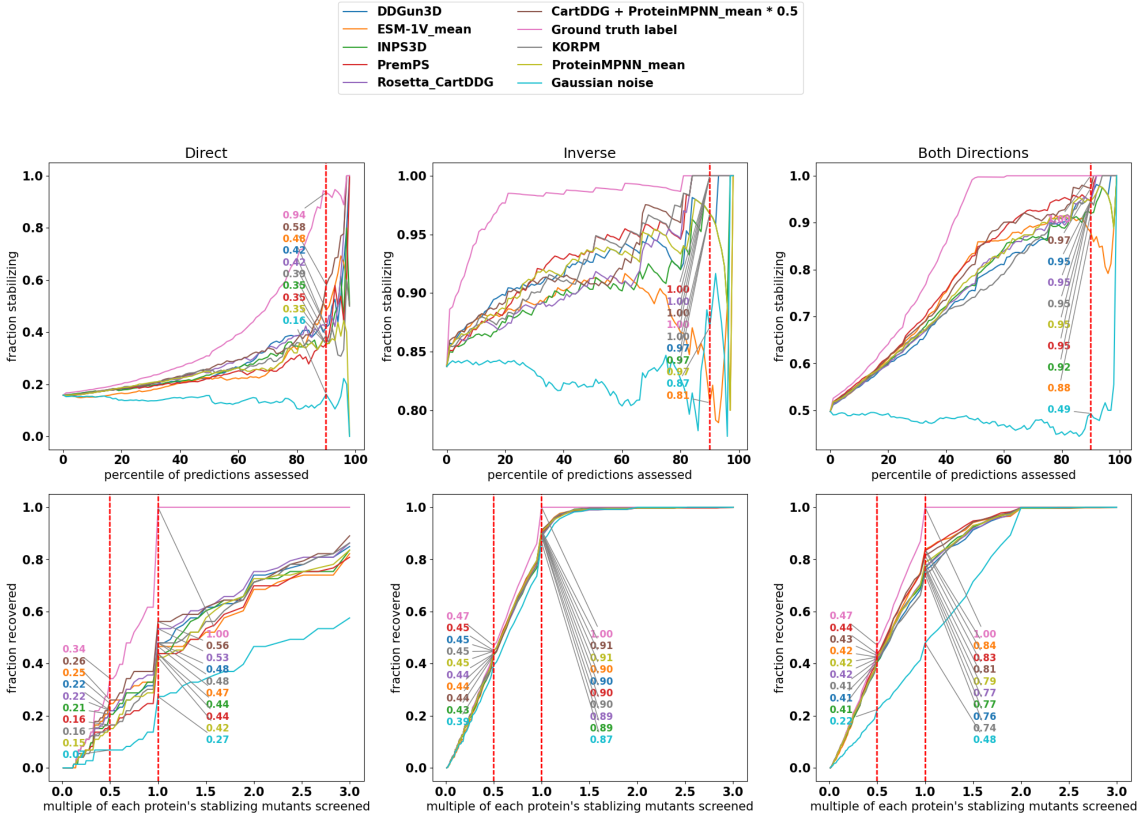


**Figure 12. Visualization of Stable Mutant Enrichment by Ensemble in S461 (All Directions)**. The top plots shows percentile-wise precision: for a given percentile of top scoring mutants according to the models, what fraction are actually stabilizing? Note that the y-axes have different ranges. In the bottom plots, the recovery (recall) of all mutations with positive stability measurements is assessed with a screening budget of various fractions of experimentally characterized stabilizing mutations per protein. There is not exactly 50% recovery at 0.5x for the ground truth labels due to rounding-down of odd numbers of stabilizing mutations. Annotations of exact scores for 0.5x, and 1.0x screening budgets are recorded.
